# The missing fraction problem as an episodes of selection problem

**DOI:** 10.1101/2023.04.27.538558

**Authors:** Elizabeth A. Mittell, Michael B. Morrissey

## Abstract

In evolutionary quantitative genetics, the missing fraction problem refers to the case where phenotypes seen later in life are biased because a non-random subset of those phenotypes are missing from the population due to prior viability selection on correlated traits. As any such missing fraction will bias our estimates of selection, and therefore, responses to selection, it is one potential explanation for the paradox of stasis seen in wild populations. The two components required for the missing fraction problem to arise are: (1) viability selection; and (2) correlation between later-life traits and those important for early-life survival. Although it is plausible that these conditions are widespread in wild populations, this problem has received very little attention since it was first discussed (Grafen 1988; Hadfield 2008). It is impossible to know what phenotypes would have been expressed later in life by individuals who died during an earlier episode of viability selection, which has probably put researchers off. Here we show that we can break the problem down into episodes of selection and recover either (a) true estimates of phenotypic selection for later-life traits, or (b) adjusted estimates of the response to selection, depending on the data available. Implementation of complex statistics should uncover how prevalent the problem may be across many existing datasets (the latter approach). Whereas overall, we hope that viewing the missing fraction problem as an episodes of selection problem increases motivation, and provides justification, for a shift in focus to directly studying early-life viability selection.

“…what a correlation with [lifetime reproductive success] or its components tells you depends on the causes of the natural variation in the character. It will not always be easy to discover those causes.”

— Grafen (1988)

“It is surprising that the problem of missing data has received so little attention given that viability selection is central to evolutionary biology.”

— Hadfield (2008)

## Introduction

Over the last century, evolutionary quantitative genetic methods have successfully predicted changes in traits in domestic settings and selection experiments (Walsh and Lynch 2018). In wild populations, these same methods often predict evolutionary change as many traits are estimated to be highly heritable (Mousseau and Roff 1987; Postma 2014) and subject to fairly strong directional selection (Kingsolver et al. 2001; Morrissey 2016). However, the predicted magnitude of evolutionary change is not often seen in the wild (Merilä, Sheldon, and Kruuk 2001); a phenomenon known as the ‘paradox of stasis’ (Hansen and Houle 2004). There is a range of potential explanations for the paradox of stasis in contemporary populations but many of these demand large amounts of data (reviewed in Pujol et al. 2018). In wild populations it is easier to generate the information that underlies the paradox - estimates of selection and heritability - than it is to obtain the information necessary to test these explanations. Therefore, we have little information on what explanations are most important for explaining mismatches between microevolutionary predictions and observations.

Studies of selection considering one or very few traits (or life stages/components of fitness) are likely to miss one of the main broad explanations for the paradox of stasis – antagonistic selection. Antagonistic selection occurs when the joint effects of genetic covariation and selection acting on multiple traits, or the same trait in a different life stage, causes the expected evolution of any given focal trait to differ from what it would be if it was considered in isolation. The missing fraction problem is one conceivable and relatively unexplored explanation for the paradox of stasis that falls within this larger class of explanations (Grafen 1988; Hadfield 2008). The missing (or invisible) fraction refers to the fraction of a population that is ‘missing’ when a trait of interest is measured due to viability selection having occurred in the population prior to measurement. Viability selection on a trait will change the distribution of any correlated traits later in life prior to their expression, relative to what they would have been in its absence. Considering that juvenile survival rates are typically fairly low, even for birds that can have relatively high juvenile survival (e.g., ∼60% in superb fairy-wrens; Hajduk et al. 2020; compared to 0.0031% in Chinook salmon; Scheuerell, Zabel, and Sandford 2009), a comparable fraction could be missing from the population for correlated later-life phenotypes. As we cannot measure the phenotypes of individuals that have died, the missing fraction may appear intractable, and, perhaps as a consequence, it has received little empirical attention.

When considering the potential impact of the missing fraction, the type of missing information and the timing of trait measurement is key – is it random who is being measured for a trait in a population at a certain life-stage and who is being missed? If individuals that are already dead at some point in the life cycle are missing not at random (MNAR) with respect to a trait that is expressed at the stage of measurement (i.e. if viability selection has occurred on a correlated trait), then inferences of selection based on alive individuals and their fitness will be incomplete (Hadfield 2008). Therefore, when phenotypes under apparent directional selection are *disproportionately* missing at a given life stage post-viability selection, our inferences of selection operating at advanced life stages may predict more evolution than should be expected, or could even predict evolution in the opposite direction to a population’s actual trajectory. A striking empirical example of the missing fraction based on an experimental study from Mojica and Kelly (2010) illustrates the problem (summarised in Box 1).

For evolutionary biologists working in wild populations it is natural to ask: what are we to do if it is fundamentally impossible to measure the later-life phenotypes of individuals that are already dead? To try and overcome this concern, we will think about the problem as a case of sequential episodes of selection that could be antagonistic in respect to one another. The theory for selection of multivariate traits across multiple intervals of selection is well developed (Arnold and Wade 1984b, 1984a; Wade and Kalisz 1989). As outlined above, prior viability selection on one (or more) correlated trait(s) will ultimately be the reason that individuals are MNAR for a focal trait at a given stage later in life (Hadfield 2008), and can therefore be considered as one episode among multiple episodes of selection throughout the life-cycle. Overall, we hope that a discussion of the missing fraction problem specifically in terms of episodes of selection theory will motivate studies that can better address it. In addition, we demonstrate that less information is required than might be expected to adjust our estimates of selection in wild populations to the effects of the missing fraction, and therefore, it could be subjected to much more empirical scrutiny with existing datasets.

### Box 1.

Empirical example of the missing fraction: part 1

An excellent example in an experimental system that illustrates the missing fraction in observational studies is provided by Mojica and Kelly (2010) using flower size in the yellow monkeyflower, *Mimulus guttatus*. In this experiment, Mojica and Kelly (2010) used seeds from three genotype classes with significantly different mean flower sizes. Importantly, each class contained high within-class genetic variation and were therefore not limited by the potential to change. By knowing the flower size of each genotype class, estimates of viability selection could be made for each class prior to flowering. The overall result was a positive correlation between flower size and fecundity at flowering (a common finding; Kingsolver et al. 2001; Sandring and Ågren 2009), and strong viability selection within the large-flowered genotype class. In other words, plants with larger flowers had greater reproductive success when measured at flowering, but a considerable proportion of the large flower individuals did not survive to that point so overall reproductive success for this genotype class was lower (Box Figure 1). Estimates of relative fitness would typically only include individuals measured at flowering and conclude that larger individuals should contribute more to the successive generation, leading to the prediction of an increase in flower size in future generations. By considering both fecundity and survival, Mojica and Kelly (2010) found that smaller genotypes were relatively fitter. The individuals that were lost due to viability selection and therefore not measured at flowering, the ‘missing fraction’, explained why there was no increase in flower size. As shown by Mojica and Kelly (2010), any study on adult traits using correlative selection analysis post-viability selection is at risk of missing this invisible fraction.

**Box Figure 1:**
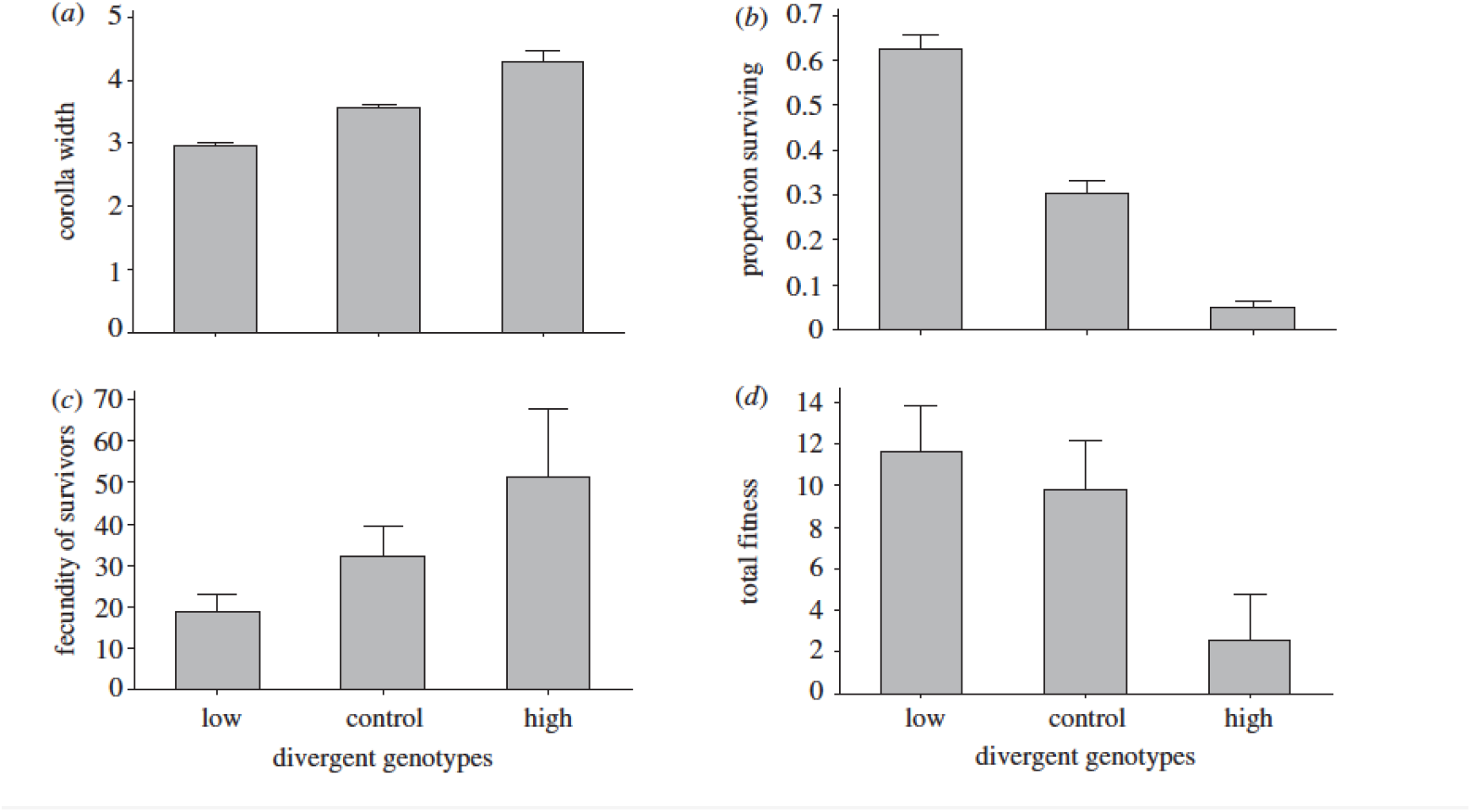
Figure reproduced from Mojica and Kelly (2010) (with permission from the Royal Society) showing the means and 95% confidence intervals for (a and c) fecundity and (b) viability measures of fitness, and (d) total fitness in the three genotype classes of *Mimulus guttatus* used in their experiment.

## Review of episodes of selection theory

Since the missing fraction problem arises from missing prior episodes of selection, we need to understand how selection combines across episodes. There are three theoretical relations and properties of modern statistical methods to know about in order to achieve this understanding.

The first key fact is that selection differentials ^1^ for any given trait are additive (Arnold and Wade 1984b). Selection differentials reflect the total selection of a trait ^2^ in a given episode, or across the entire life cycle, whether that selection is due to the effect of the trait itself on fitness, or whether the trait is correlated with a trait that directly affects fitness. If there are two relevant episodes of selection, and a trait has selection differential *S*_*i*_ in the first episode, and *S*_*ii*_ in the second, then total selection is *S*_*t*_ = *S*_*i*_ + *S*_*ii*_. The same holds for multivariate selection differentials, with the bivariate case being

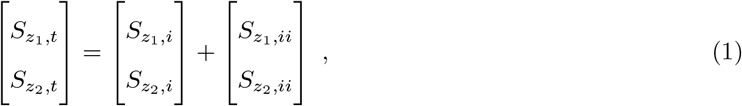

where *S*_*x,y*_ is the selection differential for trait *x* in episode *y* (e.g., 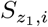 refers to the selection differential for trait 1 in episode *i*) ^3^.

The second key fact is that even if the selection differential for a focal trait cannot be directly measured, because the trait has not been expressed yet, it can be calculated if the following is known: (1) what selection has occurred on other traits; and (2) how those traits are correlated with the focal (as-yet not expressed) trait(s). If trait 1 (*z*_1_) is under viability selection at the early-life stage, then the indirect selection on the later-life trait (*z*_2_) resulting from direct selection on *z*_1_ 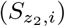 is given by the product of the phenotypic covariance of the traits 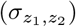 and the selection differential of *z*_1_ 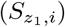 over the phenotypic variance of *z*_2_ (Equation 2). Although this relation is used for a set of traits under simultaneous selection (as in a typical selection analysis following Lande and Arnold 1983), it is directly applicable to the situation where one trait cannot have any direct effect on fitness because it has not been expressed yet. Written for a bivariate example, the contribution of selection on trait 1 to the selection differential of trait 2 is:

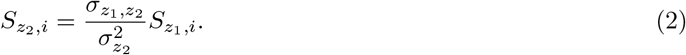

If the viability stage involves more than one trait, the standard multivariate extensions hold (Appendix 1) ^4^.

If several traits are involved in selection in episode *i*, then selection gradients for those traits can be obtained 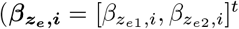, where *z*_*e*_ refers to an expressed trait), without including as-yet unexpressed traits (*z*_*n*_), as in any multivariate selection analysis. Subsequently, selection differentials of one or more non-expressed traits can be recovered with the following expression:

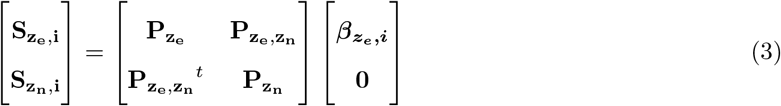

where 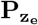 is the phenotypic covariance matrix of traits expressed and selected in the first episode (*i*), 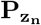 is the phenotypic covariance matrix of the traits not yet expressed in *i*, and 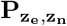 is the matrix of phenotypic covariances of expressed and selected traits with the not-yet expressed traits. All of these covariances are measured in the first episode *prior* to viability selection (see Appendix 1).

So, to calculate lifetime total selection of later-life traits, the prior selection differentials of not-yet expressed traits can be calculated by equation 3, and these differentials (that would otherwise be missing) can be combined by equation 1. It may seem problematic that these procedures require knowledge of the covariances among traits subject to prior viability selection, when that very viability selection will change those covariances before we get a chance to measure them. Fortunately, calculation of the necessary covariance is possible, and this is the third key fact for tackling the missing fraction problem: in certain circumstances, maximum likelihood techniques allow us to calculate the covariances among traits that reflect the relationships among the traits in the absence of viability selection, even if viability selection occurs (numerical example in Figure 1). The conditions for this estimation of the pre-selection covariances among traits are: (a) the traits responsible for the viability selection are measured (e.g., body size for juvenile first year over-winter survival); and (b) the missingness of traits expressed later is encoded in the data. Essentially, using the pre-viability selection phenotypes of all individuals, including those that died prior to expressing traits of interest relevant to later-life selection, and encoding their later, missing phenotypes as missing (rather than excluding their entire records), unbiased estimation of the necessary variances and covariances is possible by using either a maximum likelihood or Bayesian approach. This is because, unlike the standard moment-based estimators of variance and covariance, these approaches allow for missing data (see Figure 1 and Appendix 2 for comparison of estimated covariances).

**Figure 1:**
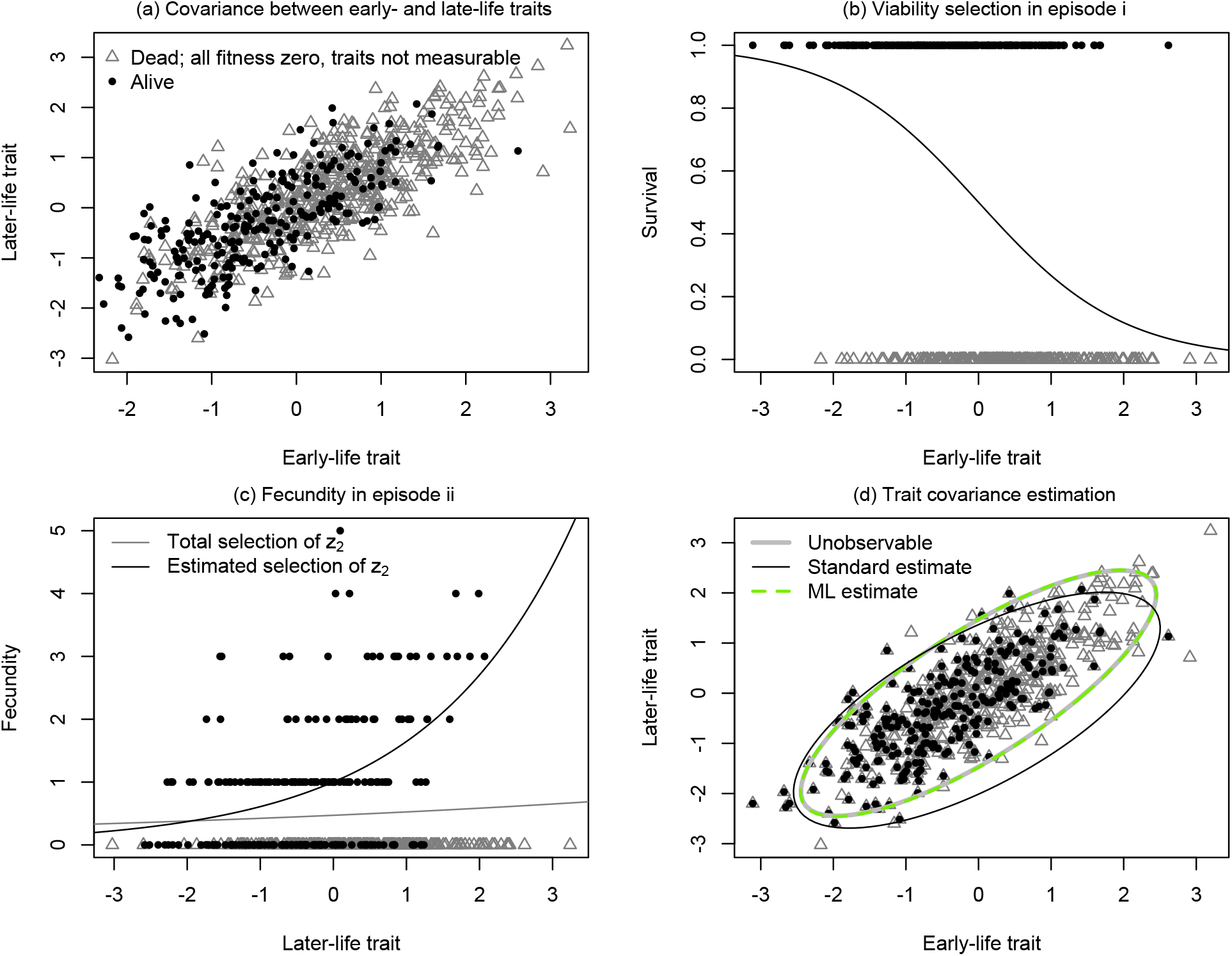
A numerical illustration of the episodes of selection. The covariance between an early- and late-life trait is shown in (a; *z*_1_ and *z*_2_, respectively). The prior viability selection of trait *z*_1_ is shown in (b). In (c) the estimated fecundity selection of *z*_2_ (from observable data) is shown in black and the true total selection on *z*_2_ is shown in grey. (d) shows the covariance of the two correlated traits when it is observable and unobservable (due to prior viability selection preventing *z*_2_ being expressed). The unobservable covariance is shown in grey, the standard estimate of covariance based on observable values is shown in black and the expectation maximum likelihood (ML) estimate of the covariance using only observed values of *z*_2_ is shown in green. Throughout individuals that died in early-life, did not express *z*_2_ and so have no fecundity, are shown in grey triangles. Individuals that survived are shown in black.

### Episodes of selection as a solution to missingness in practice

In order to demonstrate the theory outlined, simulations with simple parameters were run to compare what would be inferred about selection on a later-life phenotype using episodes of selection relative to the true values. The code for all the simulations and analyses can be found as supplementary material, as well as a comparable pedigree-based analyses on a half-sib dataset.

#### Set-up

Data for two genetically and phenotypically correlated traits were generated in 500 half-sib families each with 5 sibs according to:

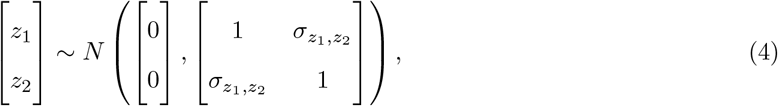

where

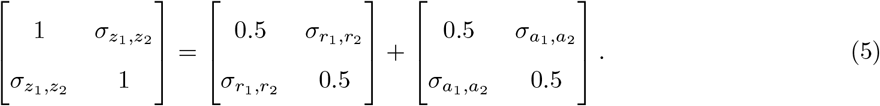

The traits were expressed early (*z*_1_) and late (*z*_2_) in life and mean trait values were set to zero. The variance-covariance matrices of total phenotype and sib-effects on the correlated, previously selected trait, and the focal trait, are shown in matrices with 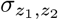 and 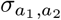 representing the phenotypic and genetic covariance between the two traits, respectively. 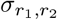 is the residual covariance. The 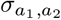 was set to 0.4, and 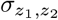 was varied from 0.8 for highly correlated traits, that are likely to cause a missing fraction problem, down to 0.4 for more weakly correlated traits.

Trait *z*_1_ affected the viability of individuals, *j*, within the population

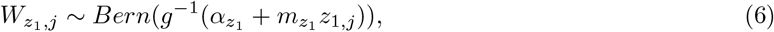

and trait *z*_2_ affected reproduction

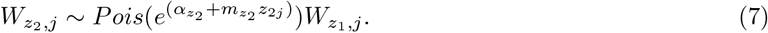

The inverse logit function (*g*^*−*1^) was used to generate trait-dependent survival probabilities 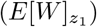 from episode *i* to *ii*. The intercepts 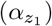 for *z*_1_ were set to zero, which means approximately 50% of the individuals survive viability selection in episode *a*. A range of 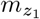 values, which generate the strength of the association between the phenotype and fitness, were used to assess the impact of the missing fraction under different viability selection regimes (an example of what these data look like for the case where 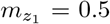 and 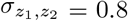 is shown in Figure 1c). The strength of fecundity selection 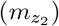 was set to 0.25 throughout. For *z*_2_the means and variances were calculated based on what would have been present in the population had all individuals survived the first episode of (viability) selection, as well as the means and variances for those individuals that survived viability selection for *z*_1_ and would have been seen by an observer (*z*_2_ in *y* = *ii*). This meant that the true 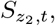 how large 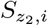 was and what would have been estimated from measurement of *z*_2_ in the second episode alone 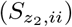 were all known. The simulation was run 1000 times for each 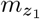.

#### Analysis

Information on the early-life trait phenotypes (*z*_1_ in *y* = *i*) was used, alongside the observed later-life phenotypes (*z*_2_ in *y* = *ii*), to obtain estimates of phenotypic covariance between the early-life and later-life traits, and the variance in the later-life trait, using expectation maximisation (a standard maximum likelihood algorithm; see Figure 1 and Appendix 2; function `emNorm` in `R` package `norm2`; R Core Team (2018); Schafer (1997)]. These estimates were combined with 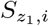 (and 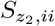for 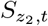) in equation 2 to obtain 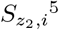. True values for an infinite population were obtained for comparison (see Appendix 3).

#### Results and conclusion

Using episodes of selection, estimates from simulated data (where later-life phenotypes of individuals that died early in life were withheld) closely matched the true values for prior viability selection of trait 2 (Figure 2) and the phenotypic covariance between trait 1 and trait 2. When the phenotypic covariance is high, using episodes of selection brings the estimates of total lifetime selection on trait 2 significantly closer to the true values for any 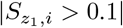; confidence intervals do not overlap 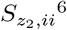 was not accounted for. When the phenotypic covariance is reduced to 0.4, 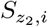 is lower and 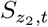*t* becomes closer to 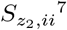. However, the 95% confidence intervals of 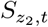 do not overlap 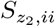 for any 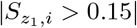, which indicates that this method gets estimates closer to true 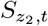 when prior viability selection is reasonably strong. These results demonstrate that it is worth investigating early-life traits believed to be important for early-life survival, and quantifying the viability selection acting upon them, alongside estimates of later episodes of selection on later-life traits. This allows identification of the causal agent (prior unseen viability selection) of selection on a focal later-life trait as well as improving predictions of evolutionary trajectories. In order to achieve this more intensive studies that focus on juvenile viability selection and collect targeted phenotypic data early in life may be required in many cases. Despite the data-demand of this approach, using it would improve our understanding of prior viability selection and its impact on the later-life traits.

**Figure 2:**
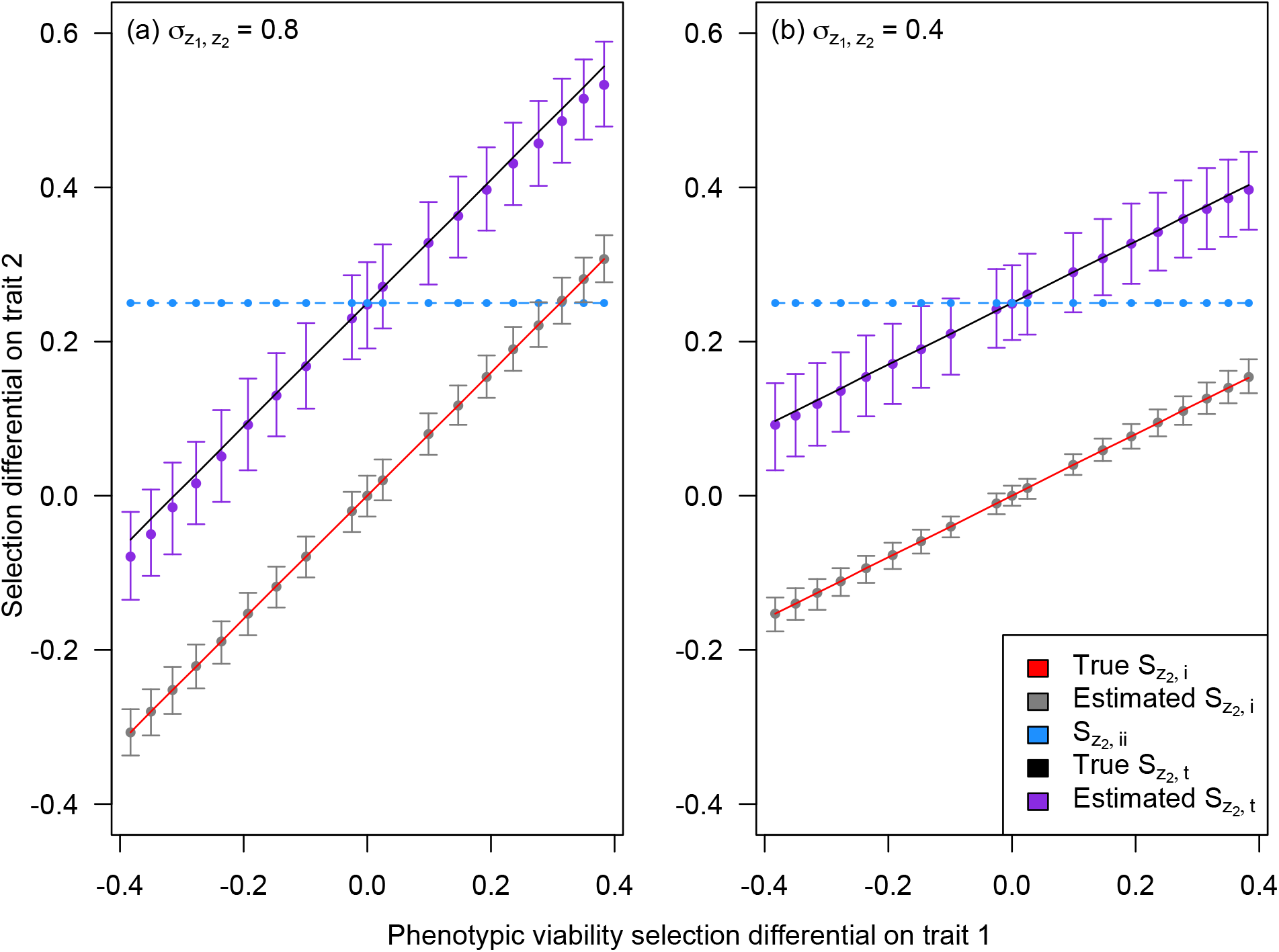
Demonstration of episodes of selection theory as a solution to missingness in later-life phenotypes due to indirect selection early in life. Data was simulated for two traits that are expressed early (trait 1) and late (trait 2) in life and have: (a) a high phenotypic correlation 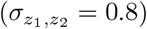 and (b) a weaker phenotypic correlation 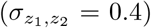. An example of what these data look like for the case where 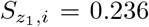 and 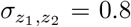 is shown in Figure 1c. The blue points represent the fecundity selection on trait 2 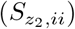. This fecundity selection is what would be estimated for total lifetime selection on trait 2 without accounting for the prior viability selection acting via trait 1. The true and estimated effect of prior viability selection on trait 2 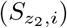 is shown in red and grey, respectively. The true and average estimated lifetime selection differential for trait 2 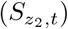 is shown black and purple, respectively. The 95% spread of estimated values are also shown.

### Genetic signatures of prior viability selection theory

Collecting data to study early life viability selection *alongside* selection on later-life traits would be the most satisfying solution for the missing fraction problem. However, when it is difficult to study early-life viability selection or we seek to analyse existing later-life data, there are approaches that can greatly improve estimates of the selection of later-life traits by reducing their susceptibility to prior viability selection on correlated traits.

The effects of the missing fraction on estimates of later-life selection can be reduced by estimating the genetic covariance between early-life survival and later-life traits. This can be done when pedigree relationships between individuals that die early, and those that go on to experience later-life periods of selection are known by utilising GLMMs for quantitative genetic analysis (e.g., Kruuk 2004; Hadfield 2010; De Villemereuil et al. 2016)^8^. The phenotypes of those that died do not need to be known, for either the later-life trait being studied or those traits that are under viability selection. Survival itself is a trait that is by definition under viability selection, and although there is no meaning to a phenotypic covariance between early-life survival and later-life traits (it is undefined), the genetic covariance is estimable and can be utilised.

From a quantitative point of view, the genetic correlation of early-life survival with later-life traits indicates when evolutionary change in a later-life trait(s) may be constrained due to antagonistic selection, which would reveal one possible cause of the paradox of stasis in wild populations. Importantly, there is also a clear quantitative meaning to the genetic covariance of early life survival with subsequently selected traits. Just like phenotypic selection differentials, genetic selection differentials, or the genetic covariances of traits with relative fitness, are additive across sequential stages of the life cycle. The genetic selection differentials (*S*_*a*_) associated with the prior viability selection of later-life traits fitness is measured by the early-life survival rate, and so

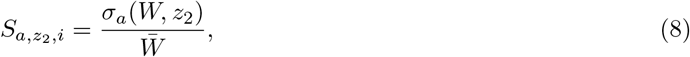

where *W* is the early-life survival rate, or absolute fitness in episode *i*^9^.

The genetic selection differential of a later-life trait, implied by a phenotypic selection analysis and estimate of genetic parameters, is the evolutionary prediction from that analysis, based on the breeder’s equation (or related equations; Lande 1979; Morrissey 2014). So, given an analysis of the genetic covariance of a later-life trait with prior survival, and an analysis of selection of that trait later in life, lifetime selection is given by

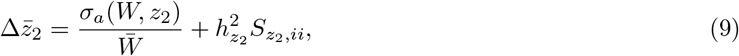

where *W* is the early-life survival rate and *h*^2^ is the narrow-sense heritability of 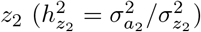.

### Genetic signatures of prior viability selection in practice

The previous section outlines how more insight can be gained into *possible* mechanisms that might be behind the missing fraction in a study by estimating genetic covariances of traits of interest in a given episode of selection with previous survival (equation 8). Doing this, genetic correlations between early-life survival and later-life traits can be found, providing an understanding of the size and effect of the missing fraction ^10^. Genetic covariances represent the association of breeding values of the trait with early-life survival. Models of the total response to selection for focal traits can then incorporate this information (equation 9), which should improve these estimates. This should be achievable for many individual-based studies where information is available about the existence of individuals that died early in life (e.g., non-hatching eggs) and their relatedness with alive individuals, as it does not require any phenotypic knowledge of those individuals that died. For this analysis the same simulated sibship data was used from the previous example ^11^.

#### Analysis

A model-based inference of the contribution of prior *genetic* viability selection to the selection differential of a later-life trait is possible without knowing early-life phenotypes when who died, and the genetic relationship between those who died and the survivors is known. The 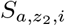 was estimated using a pedigree in an animal model framework, where the genetic component *a* is shared between early-life survival (*W*) and the later-life trait (along the lines of Steinsland et al. 2014): ^12^

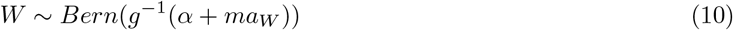

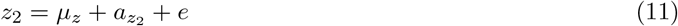

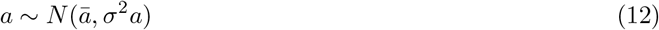

where *μ*_*z*_ and *e* are the mean and residual of *z*_2_, respectively, *ā* and 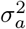 are the shared mean and variance for the genetic component of both traits, respectively, and *α* and *m* are the intercept and slope for the survival model, respectively. The model was implemented in the R package `JAGS` (Plummer 2019) using a logistic regression (*g*^*−*1^) on early-life survival. The estimates of *α, m, μ*_*z*_ and 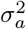 were used to calculate 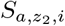 following the equations 14, 15 and 16 (outlined in Appendix 3; Morrissey 2015).^13^

#### Results and conclusion

We were able to infer the genetic prior viability selection differentials of trait 2 well (Figure 3). These results do not depend on the phenotypic covariance of the two traits (just the genetic covariance), and therefore, there is little difference to be seen between Figures 3a and 3b. Our estimates of total genetic selection of trait 2 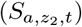 are greatly improved when 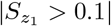. However, the distribution of estimates around 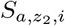 is fairly wide, and therefore, the 95% distribution of estimate 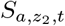 overlaps 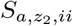 when 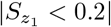. This method only requires that knowledge of individuals that died or survived early periods of viability selection, but not their phenotypes (or in any case, not the phenotypes for those traits that are under viability selection), how these individuals were related and the phenotypes of the survivors for the focal later-life trait. Therefore, it is likely that many datasets already exist where the effects of the missing fraction (if present) on estimates of selection on later-life traits could be reduced by implementing this method.

**Figure 3:**
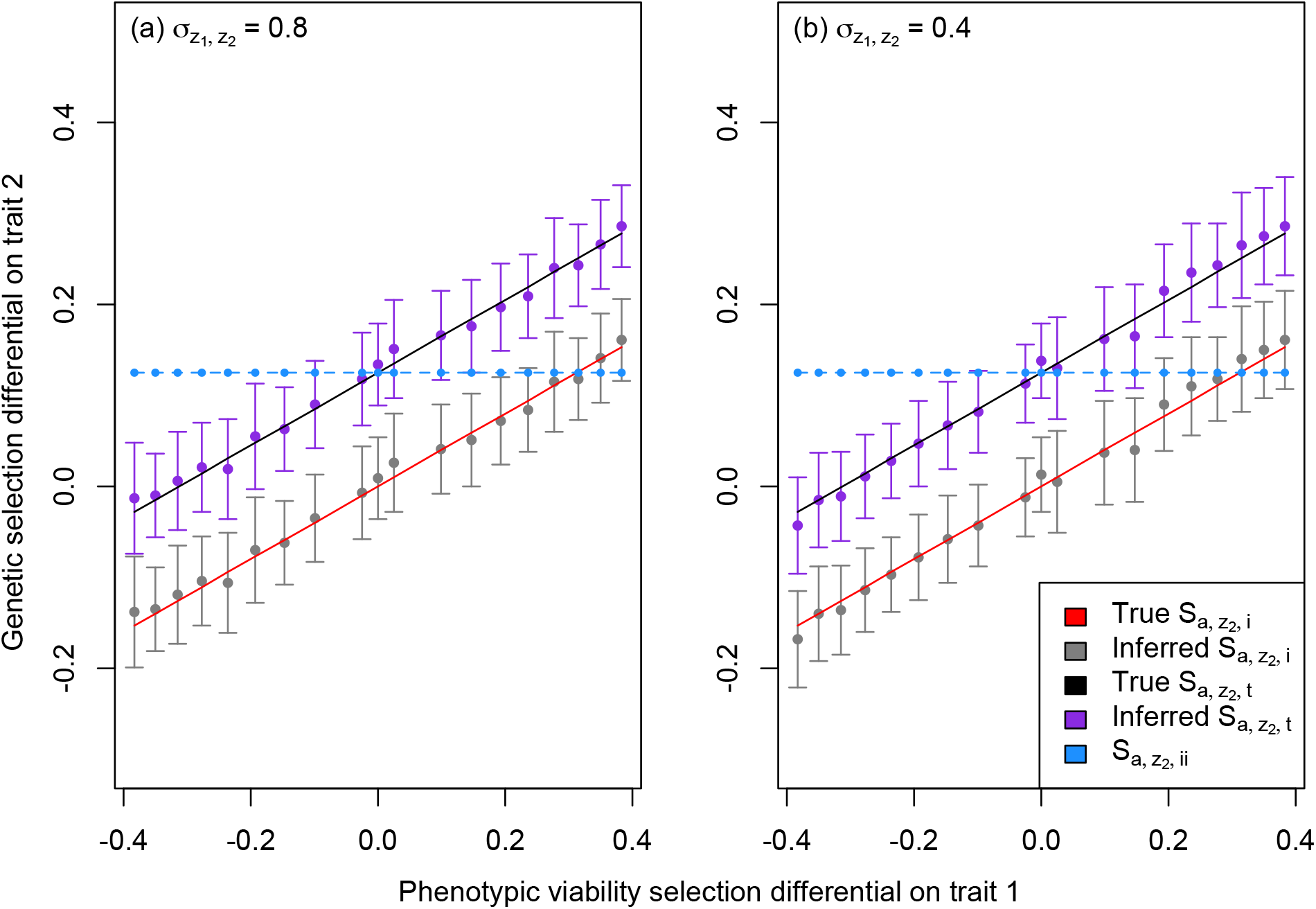
Demonstration of genetic signatures of prior viability selection as a solution to missingness in later-life phenotypes due to indirect selection early in life. Data was simulated for two traits expressed early (trait 1) and late (trait 2) in life that have (a) high phenotypic covariance 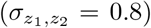 and (b) weaker phenotypic covariance 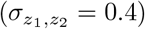. An example of what these data look like for the case where 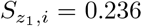 and 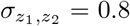 is shown in Figure 1c. The blue represents the genetic fecundity selection differential on trait 2 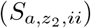. This fecundity selection is what would be estimated for total lifetime selection on trait 2 without accounting for the prior viability selection acting via trait 1. The true and estimated prior genetic viability selection differentials for trait 2 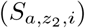 are shown in red and grey, respectively. The true and estimated lifetime genetic selection differentials for trait 2 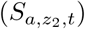 are shown in black and purple, respectively. The 95% distribution of results around the average estimated values are shown.

### A way forward: Increasing attention on prior viability selection

The theory necessary to allow empirical studies to combine information about selection throughout the life cycle into the larger picture about total selection became reasonably well developed during the 1980s (Arnold and Wade 1984b, 1984a; Wade and Kalisz 1989, 1990). However, relative to its power, this theory has been greatly under-used. There are probably various reasons for its under-use, ranging from views about how to implement selection theory and misunderstandings of what is required to do so, through to challenges in data collection and estimating relatedness in wild populations. Since individuals that do not express adult trait values will also have no reproductive fitness, it can be easy to overlook them (the missing fraction) when the focus is on the number of zygotes going into the next generation (Walsh and Morrissey 2019). It is well established that this can bias estimates of selection when this fraction is not missing at random, as demonstrated here. Increasing the focus on early-life viability selection using the theory outlined above may go some way to solve this problem.

The way that previous conceptual articles addressing the missing fraction have been framed has probably deterred research on the problem by creating a perceptual issue. Most works are written from a retrospective point of view where prior periods of viability selection were not studied and the process of data collection has been on individuals where viability selection has already occurred. These conceptual works then – correctly – give the impression that these situations are very difficult to tackle. After all, they arise from the fact that it is quite impossible to measure later-life phenotypes of individuals that have already died. However, the general situation isn’t quite so gloomy if we think about what future data collection should be motivated by the missing fraction problem. Associations between viability and subsequent traits are not mysterious - they are simply the result of viability selection at early stages in the life cycle creating data that is missing not at random. The tools to integrate this kind of selection into our view of total lifetime selection (Arnold and Wade 1984b, 1984a; Wade and Kalisz 1989, 1990) are well-established and can rely on reasonably straightforward statistical analyses (see section “Episodes of selection as a solution to missingness in practice”). We hope that by casting both the genesis and the solution to the missing fraction problem as a matter of episodes of selection increases motivation, and provides justification, for more direct inference on early-life viability selection and its incorporation into assessments of total selection. Early life viability selection is of course extremely interesting in its own right, but the fact that its study is also critical to understand total lifetime selection and thus evolution of later life traits does not so far seem to have been a clearly received message from previous thought on the missing fraction problem.

Not only could our estimates of lifetime selection be improved by comprehensively incorporating early viability selection, but it could also alleviate the paradox of stasis (Hansen and Houle 2004). Any time early-life viability selection involves a trade-off with late-life selection (Stearns 1989), not only will a missing fraction be created relative to late-life traits, but it will have a particular directionality. In particular, any such antagonistic selection would cause the association of late-life traits with lifetime reproductive success that over-estimates total lifetime selection.

An increased focus on viability selection early in life will come with certain drawbacks, but also with some very clear benefits. In many systems, early life stages are highly mobile - in many cases, early life stages are the dispersing stage. Such organisms may never be highly amenable to the study of viability selection in early life stages (although it would be impressive to tackle the problem in such systems). Another feature of early life is that early life stages are necessarily more numerous than later life stages. This may seem a drawback - any empiricist working on selection knows that keeping track of fitness components in enough individuals to estimate selection is a lot of work. However, this is also an enormous benefit. Effort intensive as they are, almost all studies of natural selection are very underpowered (Hersch and Phillips 2004; Morrissey 2016). In selection studies of adult traits, sample sizes are often limited by population size, this limitation will generally be less when working on early life traits. Therefore, more value should be placed on studies getting at early life viability selection with sufficient sample sizes to obtain reasonable precision as important contributions; even if the types of effort they require may mean that they are less multifaceted than we are often used to seeing (e.g., in relation to typical studies of sexual selection, which will often present multiple estimates of different components of reproductive success). Overall, now that the tools are more broadly accessible and it is clear that less information is required than might be expected (i.e. we do not need to know as much about the phenotypes of the missing fraction), empirical studies in wild populations, and particularly those with data across life stages and encountering the paradox of stasis, should seriously begin to consider the missing fraction.

## Conclusion

Unobservable later-life phenotypes of individuals that die due to selection early in life, and the resulting data that is missing not at random, can seriously confound studies of selection and evolution. By ignoring the dead individuals and only focusing on those alive post viability selection (grey triangles and black circles, respectively, Figure 1c), as is inherent to lifetime selection analyses of later-life traits (Grafen 1988), an estimate close to the total selection on *z*_2_ will never be obtained if there is a missing fraction problem. Although the later-life phenotypes of missing individuals will never be seen, analyses can be done that capture previous viability selection (Figure 1b). First, the theory of how early life viability selection combines with later episodes of selection to shape total selection shows that the problem is surmountable in a biologically satisfying way, and it should motivate researchers to put more effort into directly studying episodes of viability selection early in life (see section “episodes of selection as a solution to missingness”). Second, when early-life traits are not phenotyped but may be selected, there are approaches that do not require early-life phenotypic data (see section “genetic signatures of prior viability selection”). Indeed, many studies do collect early-life trait and viability data, but where this is not comprehensive, the latter approach will still be widely applicable. Overall, even if viability selection is not the main question or interest of researchers (e.g., in studies of sexual selection), it can affect estimates of selection at all stages later in life, and therefore, our understanding of the consequences of selection.

## Acknowledgements

We would like to thank Dr John K. Kelly and Dr Julius P. Mojica for sharing their work for use in Box 1, and to the Royal Society for giving permission to reproduce a figure (licence no: 1341582-1).
E.A.M was supported by a NERC research grant awarded to M.B.M (NE/R011109/1). M.B.M was supported by a University Research Fellowship from the Royal Society (London).

The authors have no conflicts of interest to declare. There is no data to archive. R scripts are in the
supplementary material.

## Appendix

### A.1 Prior viability selection explictly viewed in terms of selection gradients

The additive partitioning of selection throughout consecutive episodes of selection is less straightforward when the distribution of trait values 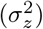 differs across episodes. Furthermore, selection gradients are typically used in natural populations where it can be difficult to discern discrete generations, and mixed models are often the preferred method to estimate selection. As selection gradients are vectors of (partial) regression coefficients of relative fitness (*w*) on trait values (*z*; Lande and Arnold 1983), they are related to selection differentials by the phenotypic variance: ***β*** = ***SP*** ^***−*1**^. This means that selection gradients are not necessarily additive over multiple episodes of selection when directional or quadratic selection occurs (Wade and Kalisz 1989). In our case the prior viability selection will have a lower weight in the linear combination of selection gradients that gives total selection, because by the time the measured episode comes around, the phenotypic variance will have been reduced and therefore the denominator will be lower. Therefore, although we note that it is likely that early life viability selection will be greater than selection in later life, it will be rare that this process will greatly change the qualitative result when selection is characterised by selection gradients instead of differentials, and we can still quantify the impact in the following manner:

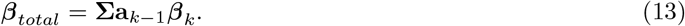

This weighting for selection gradients derived by Wade and Kalisz (1989) allows their additive partitioning, with 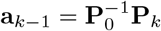. In this way, **a**_*k−*1_ weights the selection gradients at each episode (***β***_*k*_), by combining the change in phenotypic variation from the first episode (using the inverse variance-covariance matrix; 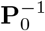) and the current episode (**P**_*k*_). N.b. if there has been no change in phenotypic variation this will be the identity matrix and therefore not give any weighting.

### A.2 Estimating phenotypic covariances in the presence of missing data

As the missing fraction occurs because of data that is missing not at random, then not only will the variance of the later-life trait change, but also the covariance between the early-life and later-life traits. If we are to use episodes of selection theory to correct our estimates of total lifetime selection on *z*_2_, then we need to be able to obtain an unbiased estimate of the covariance between *z*_1_ and *z*_2_. Standard methods of estimating covariance will not work in our case where we have data MNAR. However, we can use an approach that accounts for missing data, such as maximum likelihood based methods. For example, what follows uses the expectation maximisation algorithm from the package `norm2` (Schafer 1997), implemented in `R` (R Core Team 2018).

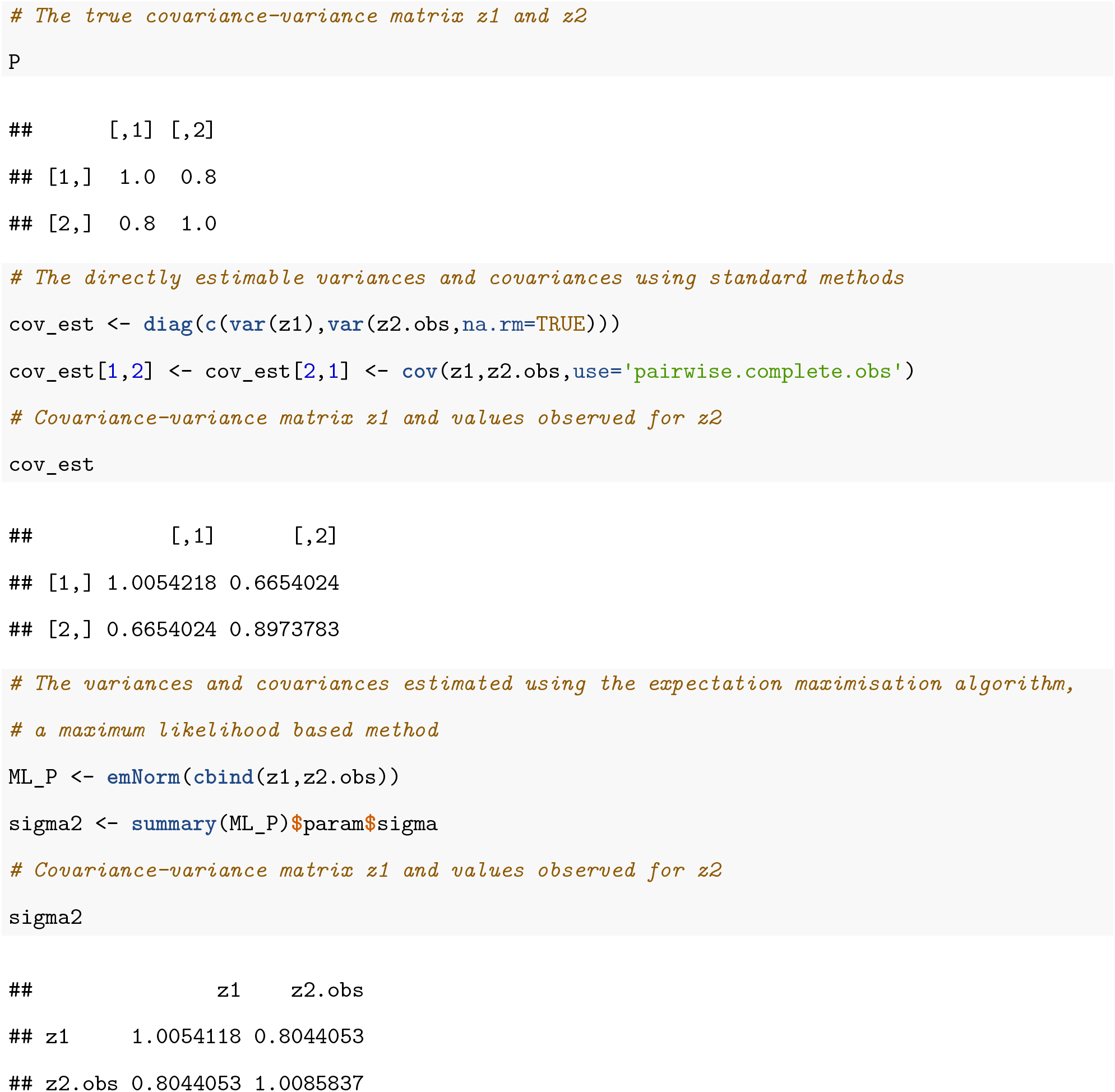

### A.3 Calculating selection differentials

True selection differentials for trait 1 in the first episode 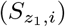 in an infinite population (section “episodes of selection as a solution to missingness in practice”) ^14^, and the estimated genetic selection differentials for trait 2 in the first 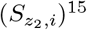 when early-life phenotypes were unknown (section “genetic signatures of prior viability selection in practice”), were calculated using integration in the equations outlined here. In the first case the true parameter values were known, whereas in the latter case estimates of these parameter values were obtained from the models as described in the main text. These parameter values were estimated from the logistic intercept (*α*) and slope (*m*), trait means (*μ*_*z*_) and standard deviation (*σ*_*z*_). We obtain mean absolute fitness in episode *i*:

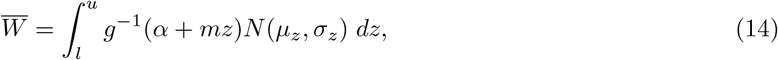

and the expectation of absolute fitness (in our case early-life survival) and the trait:

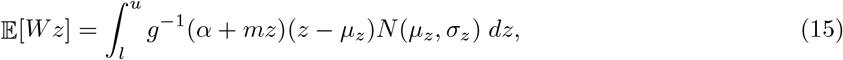

to get the prior viability selection ^16^:

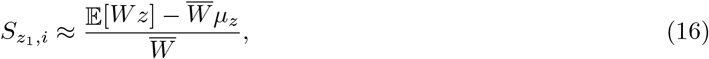

where *z* = *z*_1,*i*_; *l* = *μ*_*z*_ *−* 6*σ*_*z*_; *u* = *μ*_*z*_ + 6*σ*_*z*_.

## Footnotes

1 *S* represents a selection differential throughout. 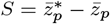 where 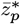 and 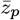 are the mean parental phenotypes that are selected (reproduce) or not, respectively.

2 Defined interchangeably as the amount that the mean phenotype changes because of selection, or by the covariance of the phenotype with relative fitness; Robertson (1966); Price (1970).

3 A corresponding relationship exists for selection gradients (Appendix 1).

4 i.e. selection gradients (*β*) are normally thought to be most relevant to a multivariate analysis, and for this we can use 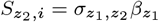 as 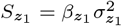.

5 These selection differentials were obtained using 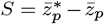 for each trait, where 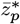 and 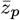 are the mean parental phenotypes that are selected (reproduce) or not, respectively.

6 This is what would be estimated as total selection on trait 2 if 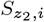

7 This is expected because 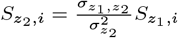.

8 Steinsland et al. (2014) carried out a similar analyses using what they refer to as a shared parameter model implemented in an alternative approximate Bayesian framework. Unfortunately, this does not seem to have been picked up on by the field.

9 Dividing through by the mean absolute fitness gives us the additive genetic covariance (*σ*_*a*_) between relative fitness assessed for early-life survival and *z*_2_. This can be calculated using equations 14, 15 and 16, where the trait of interest is a later-life trait.

10 Although, it is important to remember that using the genetic covariance between early-life survival and later-life traits does not define causal relationships at the phenotypic level that selection is acting upon, and therefore, mechanisms for the missing fraction cannot be resolved.

11 It could also be useful to point out here that when phenotypic covariance is high, the phenotypic selection differential of trait 2 early in life can also be estimated well. These results and method can be found in the supplementary materials.

12 As animal models (which require pedigree information) are used in many natural populations to partition out genetic variance from other variance components (Kruuk 2004), this should be applicable to many datasets. However, for each study system and set of questions, the correct covariates to include will need to be considered carefully by the researcher. In addition, the samples sizes will affect the accuracy of the estimates (Hersch and Phillips 2004).

13 In principle, this analysis could also be based on a bivariate response GLMM, with early-life survival and later-life traits both as response variables.

14 The true values of 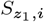 were used alongside the true phenotypic covariance and variance in trait 2 to obtain 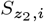.

15 i.e. the genetic covariance between early-life survival and later-life trait following equation 8

16 Here we convert the values to relative fitness as 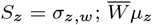 is equivalent to 𝔼 [*W*] 𝔼 [*z*].

