## Supplementary for "The missing fraction problem as an episodes of selection problem"

#### Overview

We begin this supplementary with a theoretical section on how big the missing fraction problem could be by looking at the maximum possible prior viability selection for a later-life trait. This section aims to demonstrate that even under relatively high survival rates and weaker phenotypic correlations, there is the potential for a missing fraction in our estimates of selection for later-life traits, which could be relevant for many wild populations.

Table 1 lays out the different possible levels of analyses, what data is required for each and what outcomes can be expected. For each method, episodes of selection and inference of prior viability selection, we provide code, summary data and some additional analyses for half-sib datasets, to support results using full-sibs in the main manuscript. The half-sib dataset might be more representative of study organisms where single offspring are produced at a time, and therefore, half-sibs are more likely.

Finally, we show that under certain circumstances (high phenotypic correlations between early- and late-life traits, for example, body size traits) we can actually infer prior *phenotypic* viability selection differentials for the later-life trait that are close to the true values. This doesn't require us to know what the early-life trait(s) are, just who died, the later-life phenotypes and some form of relatedness between who survived and who died (e.g., nest-of-origin or clonal line).

#### How big can the missing fraction problem be?

We are interested in how big the missing fraction problem can be because we will not always be able to obtain any information on individuals in early-life, including their existence. Two phenomena control the extent of the missing fraction problem for any later-life trait: (1) how strong viability selection was on any correlated traits earlier in life, and (2) the strength of the association between those selected traits and focal later-life traits. If we express both traits in units of their own standard deviations, then the prior viability selection of trait 2 in equation 2 of the main manuscript ($S_{z_2,i} = \sigma_{z_1,z_2}\beta_{z_1}$) can be written

$$S_{z_2,i,\sigma} = S_{z_1,\sigma}\rho_{z_1,z_2}, \tag{1}$$

where $\rho_{z_1,z_2}$ is the phenotypic *correlation* between the two traits. Without knowing values of $S_{z_1,\sigma}$ and $\rho_{z_1,z_2}$, we cannot make practical use of this expression - that is the perceived nature of the missing fraction problem. However, we can think about what limits these quantities, and therefore, what an upper limit is for (unstudied) prior viability selection of later-life traits. The upper limit on prior viability selection is set by the survival rate ($W_a$). At high survival, the opportunity for selection is low. As more mortality occurs, potentially due to selection, the opportunity for selection increases. The opportunity for viability selection, or the maximum possible selection differential, in units of standard deviations, is given by

$$i_{W_a} = \frac{S^{max}}{\sigma} = \pm \frac{f_N(q_N(1 - W_a))}{W_a}. \tag{2}$$

This is a little bit different from the general notion of the opportunity for selection, i.e., the variance in relative fitness, which is also thought of as setting an upper limit on the strength of selection. The reason is that a trait cannot have a perfect linear relationship with fitness in an episode of viability selection, because survival is composed of two classes, and therefore maximum viability selection occurs when there is a perfect step function relating a trait to survival.

Supplementary Figure 1 provides a visualisation of the upper limit of effects of prior viability selection as described by supplementary equation 2. The range of possible prior viability selection differentials is illustrated for selection on previously expressed traits with correlations of $\pm 1$ and $\pm 0.5$ (as $\rho_{z_1,z_2}$ is bound $|0-1|$) with a focal later-life trait. The ranges of possible prior viability selection are shown in relation to the latest meta-analytic estimate of the range of values typical of selection on traits in general (mainly later-life traits; Supplementary Figure 1 dashed lines; Morrissey 2016). The mortality rates that often occur early in life in natural populations (e.g., up to 70% in songbirds; Martin et al. 2018) allow for prior selection that greatly exceeds the typical strengths of selection we see for traits later in life.

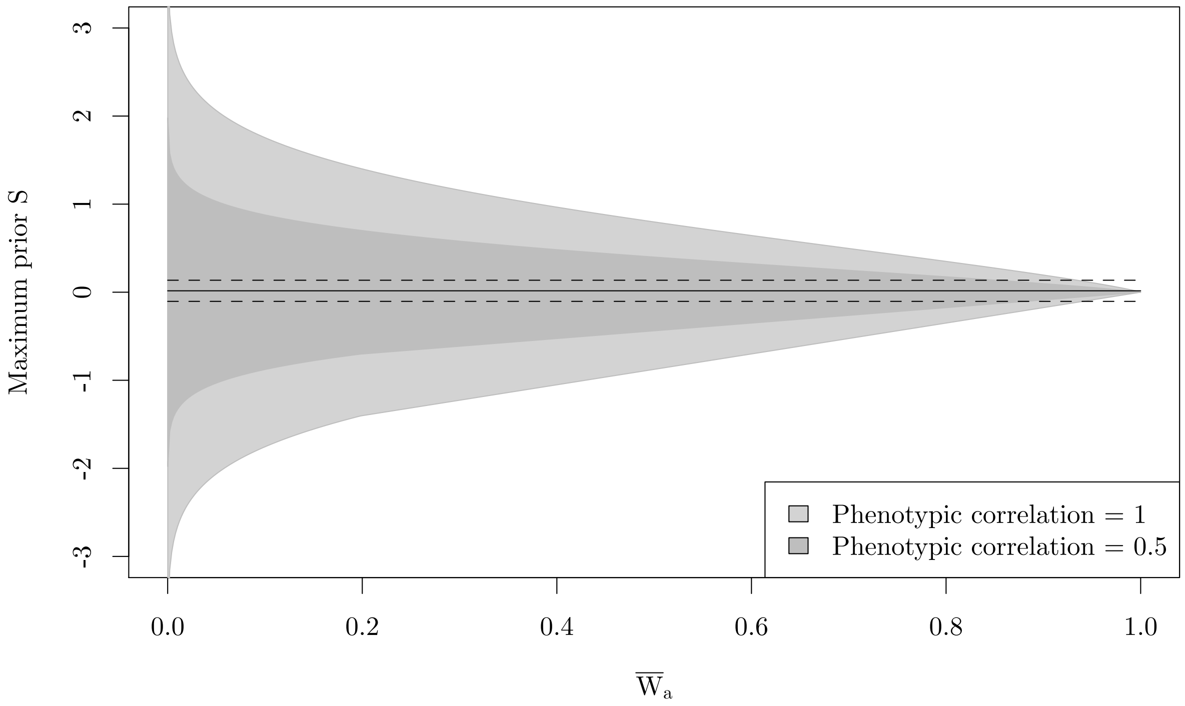

Figure 1: The upper limits of the strength of prior viability selection on an as-yet unexpressed later-life trait as a function of the survival rate, $\bar{W}_a$. This is the maximum prior viability selection that could be missing from our estimate of net selection on a focal trait based on measurements only made in later episodes of selection. Phenotypic correlations between the selected early-life trait and the as-yet unexpressed later-life trait of $\pm 1$ and $\pm 0.5$ are shown in light and dark grey, respectively. The dashed black lines show the latest range of estimates for selection on mainly later-life traits from a meta-analysis (Morrissey 2016).

However, how strong selection can be and how strong it actually is, might be very different things. Supplementary Figure 2 shows prior selection gradients that would be obtained under truncation selection, but in reality early-life survival is not likely to be so discriminating. Despite that, prior viability selection would not have to be nearly as strong as it can be in order to greatly change our impression of total selection of later-life traits. For a given prior survival rate, the other determinant of the strength of prior selection on a later-life trait is its correlation with the previously selected trait(s). These correlations could of course take almost any value, but it seems that very strong correlations could often be relevant. For example, aspects of body size are likely to be very strongly correlated across ontogeny.

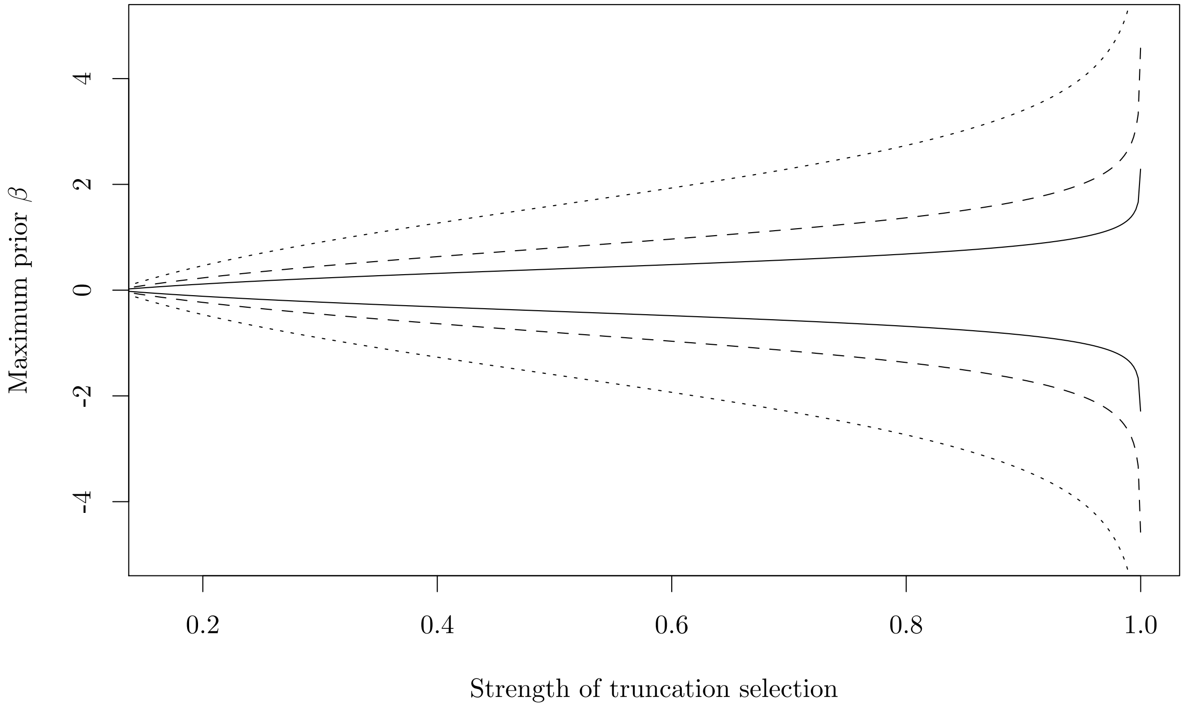

Figure 2: The maximum prior $\beta_{z_1}$ under truncation selection where $\sigma^2_{z_1} = |2|$, $\sigma^2_{z_1} = |1|$ and $\sigma^2_{z_1} = |0.5|$ are shown in solid, dashed and dotted lines, respectively.

### Levels of analyses

**Table 1:** The data requirements for each of the analyses, the general method and the outcome that can be expected.

| Analysis | Data required | Method | Outcome |
|---|---|---|---|
| Genetic signatures of prior viability selection. | (i) The existence of individuals that died early in life (e.g., non-hatching eggs).<br>(ii) The relatedness of these individuals who died with individuals who are alive.<br>(iii) The phenotype of interest of those individuals who are alive. | Estimate the genetic covariances between the trait of interest in a given episode of selection with previous survival. | Find out if genetic correlations exist, but not the underlying causal explanation, i.e. the phenotypes that viability selection is acting upon.<br>Mitigate against the missing fraction caused by prior viability selection. |
| Episodes of selection. | (i) Phenotypes of early-life traits that are under viability selection in all individuals.<br>(ii) Who survived this viability selection (to estimate the phenotypic selection differential on the early-life traits).<br>(iii) Phenotypes of later-life traits of interest for alive individuals (to estimate the covariance between the early-life and later-life traits). | Estimates of total lifetime selection corrected using information about selection in prior episodes. | Identify and quantify viability selection acting upon early-life traits that could be important for early-life survival.<br>Quantify the missing fraction and its causal agents. |
| Inference of prior *phenotypic* viability selection. | (i) The existence of individuals that died early in life (e.g., non-hatching eggs).<br>(ii) The relatedness of these individuals who died with individuals who are alive.<br>(iii) The phenotype of interest of those individuals who are alive.<br>(iv) Biological reason to believe high phenotypic correlation between early-life trait(s) responsible for early-life survival and later-life trait(s) of interest, although no measurements of early-life trait(s) are required | Estimate the phenotypic covariances between the trait of interest in a given episode of selection with previous survival. | Mitigate against the missing fraction caused by prior viability selection. |

### Episodes of selection

#### Simulated data – sibships

The scenario used for simulating the datasets within the main manuscript took relatedness information from sibships. This is the more general scenario as it does not require us to know the relatedness of all individuals within a population, or both parents, we only need to know the identity of one parent (e.g., it is common to know the mother). Therefore, it could represent parental groups where one parent is known (e.g., nest-of-origin), clonal lines or common environment effects. *This setup assumes that the offspring are full-sibs, which is a consideration when being applied to empirical datasets where nest-of-origin may contain some half-sibs depending on the study system.*

The data was simulated and these analyses were run independently 1000 times each for a range of strengths and directions of viability selection: $\beta_{z_1} = [-0.9, -0.8, -0.7, -0.6, -0.5, -0.4, -0.3, -0.2, -0.05, 0, 0.05, 0.2, 0.3, 0.4,$

0.5, 0.6, 0.7, 0.8, 0.9]. This was done for high and weaker phenotypic covariances between trait 1 and trait 2 ($\sigma(z_1, z_2) = [0.8, 0.4]$). An example summary of these data for four families/sibships can be found in Table 2.

Code for data simulation:

```
# Libraries required
library(mvtnorm)

## SIMULATION SETUP FULL SIBS ##
n.sibs <- 5 # Number of sibs per sibship (offspring of each parent)
n.sires <- 500 # Number of families (i.e. sample size, which could be number of occupied nest-boxes)
alpha <- 0 # Logistic regression intercept for z1 (early-life trait)
beta <- 0.9 # Controls the strength of viability selection on z1
# A range of beta were used in simulations:
betas <- c(-0.9,-0.8,-0.7,-0.6,-0.5,-0.4,-0.3,-0.2,-0.05,0,0.05,0.2,0.3,0.4,0.5,0.6,0.7,0.8,0.9)
S0 <- matrix(c(0.5,0.4,0.4,0.5),2,2) # Genetic var-covar matrix between z1 and z2 (later-in-life trait)
# Genetic covariance set at 0.4
P0 <- matrix(c(1,0.8,0.8,1),2,2) # Phenotypic var-covar matrix
# Phenotypic covariance set at 0.8 for highly correlated traits,
# 0.4 for more weakly correlated traits

# Generate phenotypes
d <- expand.grid(sire=1:n.sires,sib=1:n.sibs)
s <- rmvnorm(n.sires,c(0,0),S0) # Genetic component of the two traits for each family
z <- s[d$sire,] + rmvnorm(n.sires*n.sibs,c(0,0),P0-S0) # Phenotypes for each individual
# in the offspring generation
d$z1 <- z[,1] # Early-life trait
d$z2 <- z[,2] # Later-life trait

# Do individuals survive to express z2 or not?
inv.logit <- function(x){exp(x)/(1+exp(x))} # Define the inverse logit
FF1 <- function(z){rbinom(length(z),1,inv.logit(alpha+beta*z))} # Prior survival function for z1
# Generates binomial output for surviving (1) or dying (0) from early-life viability selection
# As alpha is set at 0 then 50% of the population will survive/die

# Prior survival from viability selection on z1
d$W1 <- FF1(z[,1])
# Consequence for z2
d$W2 <- rpois(length(z[,1]),exp(0+0.25*z[,2]))
# Poisson generates a range of numbers of offspring, and therefore, fitness at the later-life stage

# Total lifetime fitness
d$W <- d$W2*d$W1
# Fitness observed for z2
d$W_obs <- d$W2
# z2 has no fitness when the individual died in the first episode
d$W_obs[which(d$W1==0)] <- NA

# The phenotypes as observed in the focal episode
# (i.e. phenotypes of those already dead must be NA)
d$z2_obs <- d$z2
d$z2_obs[which(d$W1==0)] <- NA
```

Table 2: Example summary of simulated data from one run for four families. The trait values for each individual, the associated fitness, and the fitness we observe in the population are shown. In this example $\beta = 0.9$ (controls the strength of viability selection).

| Family | Offspring | Trait 1 | Trait 2 | Trait 1 fitness | Trait 2 fitness | Total fitness | Observed fitness | Expressed trait 2 |
|---|---|---|---|---|---|---|---|---|
| 1 | 1 | -0.491 | -0.947 | 1 | 1 | 1 | 1 | -0.947 |
| 1 | 2 | -0.524 | -0.063 | 0 | 2 | 0 | NA | NA |
| 1 | 3 | -0.123 | 0.271 | 1 | 0 | 0 | 0 | 0.271 |
| 1 | 4 | -0.818 | -0.711 | 0 | 2 | 0 | NA | NA |
| 1 | 5 | 0.300 | 0.278 | 0 | 2 | 0 | NA | NA |
| 2 | 1 | 1.483 | 0.372 | 1 | 0 | 0 | 0 | 0.372 |
| 2 | 2 | 1.040 | -0.666 | 0 | 2 | 0 | NA | NA |
| 2 | 3 | 0.723 | 0.118 | 1 | 1 | 1 | 1 | 0.118 |
| 2 | 4 | 0.829 | -0.020 | 1 | 1 | 1 | 1 | -0.020 |
| 2 | 5 | 0.711 | -0.481 | 0 | 3 | 0 | NA | NA |
| 3 | 1 | -1.671 | -1.446 | 0 | 0 | 0 | NA | NA |
| 3 | 2 | -0.972 | -1.235 | 0 | 1 | 0 | NA | NA |
| 3 | 3 | -1.927 | -1.335 | 0 | 0 | 0 | NA | NA |
| 3 | 4 | -1.519 | -0.593 | 0 | 1 | 0 | NA | NA |
| 3 | 5 | -0.825 | -0.439 | 0 | 1 | 0 | NA | NA |
| 4 | 1 | -0.018 | 0.203 | 0 | 2 | 0 | NA | NA |
| 4 | 2 | -1.304 | -1.355 | 0 | 3 | 0 | NA | NA |
| 4 | 3 | 0.364 | -0.258 | 1 | 0 | 0 | 0 | -0.258 |
| 4 | 4 | 0.661 | -0.106 | 1 | 1 | 1 | 1 | -0.106 |
| 4 | 5 | 0.538 | 1.067 | 1 | 2 | 2 | 2 | 1.067 |

```r
## SIMULATION RESULTS FULL SIBS ##

# True covariance-variance matrix z1, all z2 and values observed for z2
covar <- cov(d[,c("z1","z2","z2_obs")],use="pairwise.complete.obs")

# Composite function to calculate the prior viability selection differential from the
# logistic intercept and slope (a and b), trait mean and sd (m and s)
S_logistic <- function(a,b,m,s){
# get the sample range
    low <- m - 6*s; up <- m + 6*s;
# function to get absolute fitness
    fun <- function(z,a,b,m,s){(exp(a+b*z)/(1+exp(a+b*z)))*dnorm(z,m,s)}
# obtain absolute fitness
    muW <- integrate(f=fun,lower=low,upper=up,a=a,b=b,m=m,s=s)$value
# function to get covariance between absolute fitness and trait 2
    ffun <- function(z,a,b,m,s){(z-m)*(exp(a+b*z)/(1+exp(a+b*z)))*dnorm(z,m,s)}
# normalise to relative fitness
    return((integrate(f=ffun,lower=low,upper=up,a=a,b=b,m=m,s=s)$value-m*muW)/muW)
}

# Phenotypic selection on trait 1
# In an infinite population across known parameters
S1a_inf <- S_logistic(0,beta,0,sqrt(P[1,1]))
# Calculation of what prior viability selection on trait 2 should be
S2a_inf <- (S1a_inf/P[1,1])*P[1,2]

# Phenotypic S on trait 1 in simulated data
# Known if measured early-life traits and viability selection in a focal population
S1a <- weighted.mean(d$z1,d$W1)-mean(d$z1)

library(norm2)
# Variances and covariances estimated using expectation maximum likelihood
ML_P <- emNorm(cbind(d$z1,d$z2_obs))
sigma2 <- summary(ML_P)$param$sigma
# Estimated covariance-variance matrix z1 and values observed for z2
sigma2

# Estimated phenotypic prior selection differential trait 2
S2a_est <- (S1a/sigma2[1,1])*sigma2[1,2]

# Fecundity selection on trait 2
d6 <- subset(d,d$W1==1)
S2b <- weighted.mean(d6$z2,d6$W)-mean(d6$z2)

# Total selection of trait 2
S2t <- S2a_log + S2b

# Total estimated selection of trait 2
S2t_est <- S2a_est + S2b
```

**Results** The means and 95% confidence intervals for each $\beta_{z_1}$ were taken from the 1000 independent replicates. The true values (from an infinite population) were plotted against the values that would be naïvely estimated from the simulated population in Figure 2 of the main manuscript. A summary of the means for the different parameters are shown here in Supplementary Table 3.

Table 3: Summary of the results from sibship data simulated in 1000 independent runs. All values are the mean across all runs. True and estimated values for a range of strengths of selection on trait 1 (Beta z1) and phenotypic covariances (CovP) are shown. This includes viability selection for trait 1 (S1a and S1a infin refer to, the result from the simluated population and an infinite population, respectively, with the same notation used throughout), viability (S2a), fecundity (S2b) and total (S2t) selection of trait 2, covariance between the two traits (covp z1 z2), and, variance of trait 2 (var z2).

| True_covp_z1_z2 | Est_covp_z1_z2 | True_var_z2 | Est_var_z2 | S1a_pop | S1a_infin | S2_pop | S2a_infin | S2b | S2t_pop | S2a_est | S2t_est | S2t_log | CovP | Beta_z1 |
|---|---|---|---|---|---|---|---|---|---|---|---|---|---|---|
| 0.398 | 0.398 | 0.997 | 0.996 | -0.383 | -0.383 | -0.153 | -0.153 | 0.242 | 0.089 | -0.153 | 0.089 | 0.089 | 0.4 | -0.90 |
| 0.598 | 0.597 | 0.998 | 0.995 | -0.383 | -0.383 | -0.229 | -0.230 | 0.236 | 0.007 | -0.230 | 0.006 | 0.006 | 0.6 | -0.90 |
| 0.800 | 0.801 | 1.000 | 1.002 | -0.382 | -0.383 | -0.306 | -0.307 | 0.227 | -0.079 | -0.306 | -0.078 | -0.080 | 0.8 | -0.90 |
| 0.398 | 0.398 | 0.998 | 0.997 | -0.350 | -0.350 | -0.140 | -0.140 | 0.245 | 0.105 | -0.140 | 0.105 | 0.105 | 0.4 | -0.80 |
| 0.600 | 0.599 | 1.001 | 1.000 | -0.350 | -0.350 | -0.210 | -0.210 | 0.240 | 0.029 | -0.210 | 0.030 | 0.030 | 0.6 | -0.80 |
| 0.800 | 0.799 | 1.000 | 0.998 | -0.351 | -0.350 | -0.280 | -0.280 | 0.230 | -0.050 | -0.281 | -0.051 | -0.050 | 0.8 | -0.80 |
| 0.399 | 0.400 | 1.000 | 0.999 | -0.315 | -0.315 | -0.125 | -0.126 | 0.247 | 0.122 | -0.126 | 0.121 | 0.121 | 0.4 | -0.70 |
| 0.597 | 0.597 | 0.997 | 0.996 | -0.315 | -0.315 | -0.188 | -0.189 | 0.240 | 0.052 | -0.189 | 0.051 | 0.051 | 0.6 | -0.70 |
| 0.799 | 0.798 | 0.998 | 0.997 | -0.314 | -0.315 | -0.251 | -0.252 | 0.234 | -0.017 | -0.251 | -0.018 | -0.018 | 0.8 | -0.70 |
| 0.399 | 0.400 | 0.999 | 1.000 | -0.277 | -0.277 | -0.111 | -0.111 | 0.247 | 0.135 | -0.111 | 0.136 | 0.136 | 0.4 | -0.60 |
| 0.598 | 0.597 | 0.997 | 0.996 | -0.277 | -0.277 | -0.166 | -0.166 | 0.241 | 0.075 | -0.166 | 0.075 | 0.075 | 0.6 | -0.60 |
| 0.799 | 0.799 | 0.999 | 0.998 | -0.277 | -0.277 | -0.222 | -0.222 | 0.237 | 0.015 | -0.222 | 0.015 | 0.015 | 0.8 | -0.60 |
| 0.399 | 0.398 | 0.998 | 0.996 | -0.235 | -0.236 | -0.094 | -0.094 | 0.248 | 0.154 | -0.094 | 0.154 | 0.154 | 0.4 | -0.50 |
| 0.599 | 0.600 | 0.999 | 0.999 | -0.236 | -0.236 | -0.142 | -0.142 | 0.243 | 0.102 | -0.142 | 0.102 | 0.101 | 0.6 | -0.50 |
| 0.799 | 0.798 | 0.999 | 0.997 | -0.236 | -0.236 | -0.189 | -0.189 | 0.239 | 0.050 | -0.189 | 0.051 | 0.050 | 0.8 | -0.50 |
| 0.401 | 0.400 | 0.999 | 0.998 | -0.193 | -0.193 | -0.077 | -0.077 | 0.247 | 0.170 | -0.077 | 0.170 | 0.170 | 0.4 | -0.40 |
| 0.598 | 0.598 | 0.997 | 0.998 | -0.192 | -0.193 | -0.114 | -0.116 | 0.247 | 0.132 | -0.115 | 0.132 | 0.131 | 0.6 | -0.40 |
| 0.798 | 0.798 | 0.998 | 0.997 | -0.192 | -0.193 | -0.153 | -0.154 | 0.243 | 0.090 | -0.154 | 0.089 | 0.089 | 0.8 | -0.40 |
| 0.399 | 0.400 | 0.998 | 0.998 | -0.146 | -0.147 | -0.059 | -0.059 | 0.248 | 0.189 | -0.059 | 0.190 | 0.189 | 0.4 | -0.30 |
| 0.597 | 0.597 | 0.999 | 0.999 | -0.146 | -0.147 | -0.088 | -0.088 | 0.249 | 0.160 | -0.088 | 0.161 | 0.161 | 0.6 | -0.30 |
| 0.801 | 0.800 | 1.000 | 0.999 | -0.147 | -0.147 | -0.118 | -0.117 | 0.248 | 0.130 | -0.118 | 0.130 | 0.131 | 0.8 | -0.30 |
| 0.399 | 0.398 | 0.999 | 0.998 | -0.099 | -0.099 | -0.040 | -0.040 | 0.248 | 0.209 | -0.040 | 0.209 | 0.208 | 0.4 | -0.20 |
| 0.600 | 0.600 | 0.999 | 0.998 | -0.100 | -0.099 | -0.059 | -0.059 | 0.248 | 0.188 | -0.060 | 0.188 | 0.189 | 0.6 | -0.20 |
| 0.801 | 0.801 | 1.001 | 1.001 | -0.099 | -0.099 | -0.079 | -0.079 | 0.249 | 0.170 | -0.079 | 0.170 | 0.170 | 0.8 | -0.20 |
| 0.399 | 0.399 | 0.998 | 0.997 | -0.025 | -0.025 | -0.010 | -0.010 | 0.248 | 0.238 | -0.010 | 0.238 | 0.238 | 0.4 | -0.05 |
| 0.600 | 0.601 | 0.999 | 1.001 | -0.025 | -0.025 | -0.016 | -0.015 | 0.249 | 0.233 | -0.015 | 0.234 | 0.234 | 0.6 | -0.05 |
| 0.799 | 0.799 | 1.000 | 1.000 | -0.026 | -0.025 | -0.020 | -0.020 | 0.250 | 0.230 | -0.020 | 0.230 | 0.230 | 0.8 | -0.05 |
| 0.399 | 0.398 | 0.997 | 0.997 | 0.000 | 0.000 | 0.000 | 0.000 | 0.249 | 0.249 | 0.000 | 0.249 | 0.249 | 0.4 | 0.00 |
| 0.602 | 0.601 | 1.000 | 1.000 | 0.001 | 0.000 | 0.000 | 0.000 | 0.251 | 0.251 | 0.000 | 0.252 | 0.251 | 0.6 | 0.00 |
| 0.798 | 0.798 | 0.999 | 0.999 | -0.001 | 0.000 | -0.001 | 0.000 | 0.248 | 0.247 | -0.001 | 0.247 | 0.248 | 0.8 | 0.00 |
| 0.397 | 0.396 | 0.996 | 0.996 | 0.025 | 0.025 | 0.010 | 0.010 | 0.251 | 0.261 | 0.010 | 0.261 | 0.261 | 0.4 | 0.05 |
| 0.598 | 0.597 | 0.998 | 0.997 | 0.024 | 0.025 | 0.014 | 0.015 | 0.248 | 0.262 | 0.014 | 0.262 | 0.263 | 0.6 | 0.05 |
| 0.799 | 0.799 | 0.999 | 0.998 | 0.025 | 0.025 | 0.020 | 0.020 | 0.249 | 0.270 | 0.020 | 0.270 | 0.269 | 0.8 | 0.05 |
| 0.400 | 0.399 | 1.000 | 1.001 | 0.099 | 0.099 | 0.040 | 0.040 | 0.250 | 0.290 | 0.040 | 0.290 | 0.290 | 0.4 | 0.20 |
| 0.600 | 0.600 | 1.000 | 0.999 | 0.098 | 0.099 | 0.059 | 0.059 | 0.248 | 0.307 | 0.059 | 0.307 | 0.307 | 0.6 | 0.20 |
| 0.797 | 0.796 | 0.997 | 0.996 | 0.099 | 0.099 | 0.079 | 0.079 | 0.247 | 0.326 | 0.079 | 0.326 | 0.326 | 0.8 | 0.20 |
| 0.398 | 0.399 | 0.999 | 1.000 | 0.146 | 0.147 | 0.058 | 0.059 | 0.248 | 0.306 | 0.058 | 0.306 | 0.307 | 0.4 | 0.30 |
| 0.597 | 0.597 | 0.997 | 0.996 | 0.147 | 0.147 | 0.088 | 0.088 | 0.247 | 0.335 | 0.088 | 0.335 | 0.335 | 0.6 | 0.30 |
| 0.798 | 0.798 | 0.998 | 0.999 | 0.147 | 0.147 | 0.118 | 0.117 | 0.244 | 0.362 | 0.118 | 0.362 | 0.361 | 0.8 | 0.30 |
| 0.400 | 0.400 | 1.000 | 0.999 | 0.192 | 0.193 | 0.076 | 0.077 | 0.249 | 0.325 | 0.077 | 0.326 | 0.326 | 0.4 | 0.40 |
| 0.601 | 0.601 | 1.002 | 1.002 | 0.192 | 0.193 | 0.115 | 0.116 | 0.248 | 0.363 | 0.115 | 0.363 | 0.364 | 0.6 | 0.40 |
| 0.799 | 0.798 | 0.998 | 0.997 | 0.192 | 0.193 | 0.153 | 0.154 | 0.245 | 0.398 | 0.154 | 0.399 | 0.399 | 0.8 | 0.40 |
| 0.399 | 0.399 | 0.998 | 0.998 | 0.237 | 0.236 | 0.094 | 0.094 | 0.248 | 0.342 | 0.095 | 0.343 | 0.342 | 0.4 | 0.50 |
| 0.601 | 0.601 | 1.001 | 1.001 | 0.236 | 0.236 | 0.142 | 0.142 | 0.244 | 0.386 | 0.142 | 0.386 | 0.386 | 0.6 | 0.50 |
| 0.800 | 0.799 | 0.999 | 0.999 | 0.235 | 0.236 | 0.188 | 0.189 | 0.242 | 0.430 | 0.188 | 0.430 | 0.431 | 0.8 | 0.50 |
| 0.398 | 0.399 | 0.998 | 0.997 | 0.277 | 0.277 | 0.110 | 0.111 | 0.246 | 0.356 | 0.111 | 0.357 | 0.357 | 0.4 | 0.60 |
| 0.599 | 0.599 | 0.997 | 0.997 | 0.277 | 0.277 | 0.166 | 0.166 | 0.243 | 0.408 | 0.167 | 0.409 | 0.409 | 0.6 | 0.60 |
| 0.800 | 0.799 | 1.000 | 1.000 | 0.276 | 0.277 | 0.221 | 0.222 | 0.237 | 0.458 | 0.221 | 0.458 | 0.459 | 0.8 | 0.60 |
| 0.399 | 0.398 | 0.998 | 0.997 | 0.316 | 0.315 | 0.126 | 0.126 | 0.245 | 0.371 | 0.126 | 0.371 | 0.371 | 0.4 | 0.70 |
| 0.598 | 0.599 | 0.999 | 0.998 | 0.314 | 0.315 | 0.187 | 0.189 | 0.241 | 0.428 | 0.188 | 0.430 | 0.430 | 0.6 | 0.70 |
| 0.801 | 0.800 | 1.001 | 1.001 | 0.315 | 0.315 | 0.252 | 0.252 | 0.234 | 0.486 | 0.252 | 0.486 | 0.486 | 0.8 | 0.70 |
| 0.400 | 0.400 | 1.001 | 1.002 | 0.350 | 0.350 | 0.139 | 0.140 | 0.244 | 0.384 | 0.140 | 0.384 | 0.384 | 0.4 | 0.80 |
| 0.600 | 0.600 | 1.000 | 0.999 | 0.350 | 0.350 | 0.210 | 0.210 | 0.238 | 0.448 | 0.211 | 0.448 | 0.448 | 0.6 | 0.80 |
| 0.798 | 0.798 | 0.997 | 0.998 | 0.350 | 0.350 | 0.279 | 0.280 | 0.228 | 0.507 | 0.280 | 0.508 | 0.508 | 0.8 | 0.80 |
| 0.400 | 0.399 | 0.999 | 0.998 | 0.383 | 0.383 | 0.153 | 0.153 | 0.244 | 0.397 | 0.153 | 0.397 | 0.397 | 0.4 | 0.90 |
| 0.601 | 0.600 | 1.000 | 0.998 | 0.384 | 0.383 | 0.230 | 0.230 | 0.235 | 0.465 | 0.231 | 0.466 | 0.465 | 0.6 | 0.90 |
| 0.800 | 0.800 | 1.000 | 0.999 | 0.383 | 0.383 | 0.306 | 0.307 | 0.225 | 0.531 | 0.306 | 0.532 | 0.532 | 0.8 | 0.90 |

#### Simulated data – half-sibs

In cases where there is sequential monogamy or polygamy, it may be more likely that offspring are in fact half-sibs rather than full-sibs. Below is the set-up for this simulated data. The results show that these genetic changes do not make a difference to the results using episodes of selection (Supplementary Figure 3).

```
## SIMULATION SETUP HALF SIBS ##
cvr <- 0.4
beta <- 0.9
n.sires <- 500
n.offspring <- 5

d <- expand.grid(sire=1:n.sires,o=1:n.offspring)
d <- d[order(d$sire),]
d$sire <- paste("S",d$sire,sep="")
d$dam <- paste("D",1:(n.sires*n.offspring),sep="")
d$id <- paste("O",1:(n.sires*n.offspring),sep="")

# Organise the data into a pedigree, add dam and sire records before offspring
pedigree <- d[,c("id","sire","dam")]
dam.p <- data.frame(id=unique(d$dam),sire=NA,dam=NA)
pedigree <- rbind(dam.p,pedigree)
sire.p <- data.frame(id=unique(d$sire),sire=NA,dam=NA)
pedigree <- rbind(sire.p,pedigree)

# Genetic covariance 0.4
G <- matrix(c(0.5,0.4,0.4,0.5),2,2)

# Residual variance-covariance matrix -> phenotypic covariance trait 2 0.4(G) + 0.4(R) = 0.8(P)
R <- matrix(c(0.5,cvr,cvr,0.5),2,2)

P <- G+R

# Breeding values for each trait
a <- rbv(pedigree, G)
a1 <- a[3001:5500,]
d$a1 <- a1[,1]
d$a2 <- a1[,2]

# Check 1
a[3101:3110,]
d[101:110,]

# Check 2: offspring on single parent regression of a should be 0.5 (so roughly 1 if we multiple by 2)
d$father_a1 <- a[match(d$sire,dimnames(a)[[1]]),1]
d$mother_a1 <- a[match(d$dam,dimnames(a)[[1]]),1]
summary(lm(a1~father_a1,data=d))$coefficients[2,]
coef(lm(a1~father_a1,data=d))[2]*2
summary(lm(a1~mother_a1,data=d))$coefficients[2,]
coef(lm(a1~mother_a1,data=d))[2]*2

# Residual effects
r <- rmvnorm(n.sires*n.offspring,c(0,0),R)
rownames(r) <- d$id # Add id
d$r1 <- r[,1]
d$r2 <- r[,2]

# Compose phenotypes
d$z1 <- d$a1 + d$r1
```

```r
d$z2 <- d$a2 + d$r2

# Survival
inv.logit <- function(x){exp(x)/(1+exp(x))}
surv_func <- function(x,a=0,b=beta){inv.logit(a+b*x)}
d$W1 <- rbinom(dim(d)[1],1,surv_func(d$z1)) # Fitness at viability stage

# Later-life trait
d$z2_obs <- d$z2
d$z2_obs[which(d$W1==0)] <- NA # Don't observe later-life trait for those individuals that do not survive viabilit

# Consequence for trait 2
d$W2 <- rpois(length(d$z1),exp(0+0.25*d$z2)) # poisson because want to generate a range of numbers of offspring
# Total lifetime fitness
d$W <- d$W2*d$W1
# Fitness observed for trait 2
d$W_obs <- d$W

# Format data
d$animal <- d$id
str(d)
data <- d[,c("id","W1","z2_obs","z2","animal")]
data$id <- as.factor(data$id)
data$animal <- as.factor(data$animal)
data$W1 <- as.integer(data$W1)
data <- as.data.frame(data)
str(data)
str(pedigree)
```

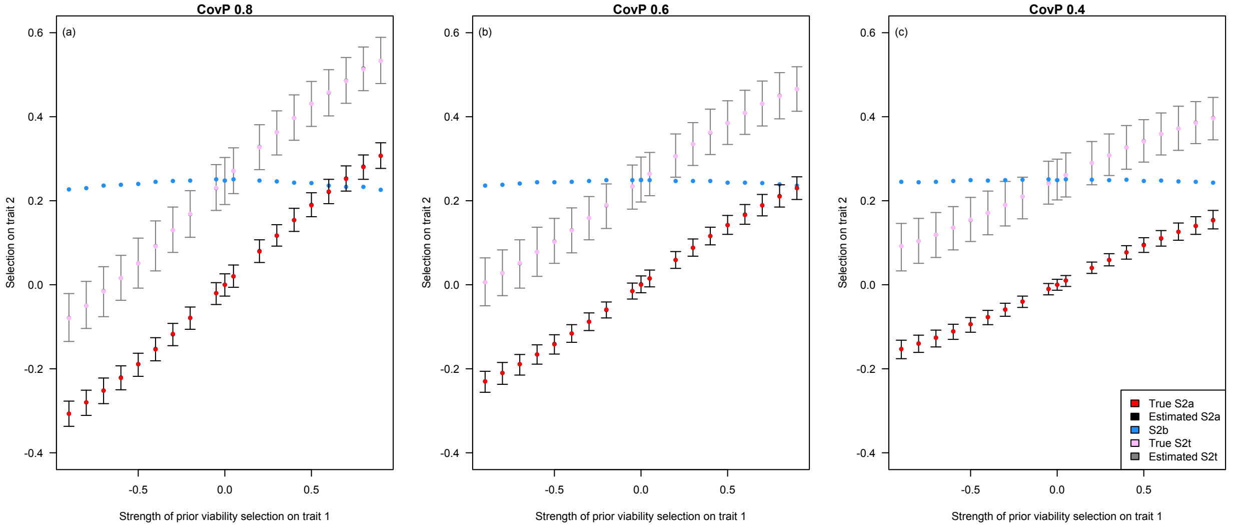

Figure 3: Results from simulated data of two traits that are expressed early (trait 1) and late (trait 2) in life and have: (a) a high phenotypic correlation ($\sigma(z_1, z_2) = 0.8$), (b) a moderate phenotypic correlation ($\sigma(z_1, z_2) = 0.6$), and (c) a weaker phenotypic correlation ($\sigma(z_1, z_2) = 0.4$). The blue points represent the fecundity selection on trait 2 ($S_{z_2,b}$). This fecundity selection is what would be estimated for total lifetime selection on trait 2 without accounting for the prior viability selection acting via trait 1. The true and estimated effect of prior viability selection on trait 2 ($S_{z_2,a}$) is shown in red and black, respectively. The true and estimated lifetime selection differential for trait 2 ($S_{z_2,t}$) is shown pink and grey, respectively. The 95% confidence intervals for the estimated values are also shown.

#### Genetic signatures of prior viability selection

##### JAGS

Below contains the few tweeks to the code above so that a pedigree-based model could be implemented to obtain an estimate of the genetic selection differential for trait 2 in the first episode ($S_{a,z_2,i}$). This is the data that the results from the manuscript are based on. *We should be aware that full-sib analyses cannot disentangle additive genetic variance from other forms of genetic variance, such as dominance variance.* The same tweeks were applied to the simulated half-sib data to allow the estimation of $S_{a,z_2,i}$. As we saw in the episodes of selection analyses, the inference method provides similar results using the half-sib data compared to the full-sib data.

```
# Libraries required
library(mvtnorm)

## SIMULATION SETUP SIRES ##
beta <- 0.7
n.sires <- 500
n.offspring <- 5
cvr <- 0.4
d <- expand.grid(sire=1:n.sires,o=1:n.offspring)
d <- d[order(d$sire),]
d$sire <- paste("S",d$sire,sep="")
d$dam <- paste("D",1:(n.sires),sep="") # Full sibs so all have same mother and father
d$id <- paste("O",1:(n.sires*n.offspring),sep="")

# Organise the data into a pedigree, add dam and sire records before offspring
pedigree <- d[,c("id","sire","dam")]
dam.p <- data.frame(id=unique(d$dam),sire=NA,dam=NA)
pedigree <- rbind(dam.p,pedigree)
sire.p <- data.frame(id=unique(d$sire),sire=NA,dam=NA)
pedigree <- rbind(sire.p,pedigree)

# Genetic covariance 0.4
G <- matrix(c(0.5,0.4,0.4,0.5),2,2)

# Residual variance-covariance matrix -> phenotypic covariance trait 2 0.4(G) + 0.4(R) = 0.8(P)
R <- matrix(c(0.5,cvr,cvr,0.5),2,2)

P <- G+R

# Breeding values for each trait
a <- rbv(pedigree, G)
a1 <- a[1001:3500,]
d$a1 <- a1[,1]
d$a2 <- a1[,2]

# Check 1
a[1001:1010,]
d[1:10,]

# Check 2: offspring on single parent regression of a should be 0.5 (so roughly 1 if we multiple by 2)
d$father_a1 <- a[match(d$sire,dimnames(a)[[1]]),1]
d$mother_a1 <- a[match(d$dam,dimnames(a)[[1]]),1]
summary(lm(a1~father_a1,data=d))$coefficients[2,]
coef(lm(a1~father_a1,data=d))[2]*2
summary(lm(a1~mother_a1,data=d))$coefficients[2,]
coef(lm(a1~mother_a1,data=d))[2]*2
```

```
# Residual effects
r <- rmvnorm(n.sires*n.offspring,c(0,0),R)
rownames(r) <- d$id # Add id
d$r1 <- r[,1]
d$r2 <- r[,2]

# Compose phenotypes
d$z1 <- d$a1 + d$r1
d$z2 <- d$a2 + d$r2

# Survival
inv.logit <- function(x){exp(x)/(1+exp(x))}
surv_func <- function(x,a=0,b=beta){inv.logit(a+b*x)}
d$W1 <- rbinom(dim(d)[1],1,surv_func(d$z1)) # Fitness at viability stage

# Later-life trait
d$z2_obs <- d$z2
d$z2_obs[which(d$W1==0)] <- NA # Don't observe later-life trait for those individuals that do not survive viability

# Consequence for trait 2
d$W2 <- rpois(length(d$z1),exp(0+0.25*d$z2)) # poisson because want to generate a range of numbers of offspring
# Total lifetime fitness
d$W <- d$W2*d$W1
# Fitness observed for trait 2
d$W_obs <- d$W

# Format data
d$animal <- d$id
str(d)
data <- d[,c("id","W1","z2_obs","z2","animal")]
data$id <- as.factor(data$id)
data$animal <- as.factor(data$animal)
data$W1 <- as.integer(data$W1)
data <- as.data.frame(data)
str(data)
str(pedigree)
```

The JAGS model code to estimate the parameters to put into the same S_logistic function as before:

```
# Jags simulation model -- genetic logistic
missingModgen <- "model{
# Priors
# Genetic
sd_a ~ dunif(0,3)
tau_a <- 1/sd_a^2
# Residual
res_sd ~ dunif(0,3)
res_tau <- 1/res_sd^2

mu ~ dnorm(0,0.0001)
alpha ~ dnorm(0,0.0001)
beta ~ dnorm(0,0.0001)

# Parents breeding values
for(j in 1:n_s){
    a_s[j] ~ dnorm(0,tau_a)
}
for(j in 1:n_d){
```

```
    a_d[j] ~ dnorm(0,tau_a)
}

# Simulation
for(i in 1:n_ind){
    # Genotypes
    a[i] ~ dnorm((a_d[dam[i]]+a_s[sire[i]])/2,tau_a*2) # Full sibs
    # Phenotypes
    z[i] ~ dnorm(mu + a[i],res_tau)
    # Observed phenotypes with little error
    z_obs[i] ~ dnorm(z[i], 1000)
# Survival probability based on the logistic fitness function
    s[i] ~ dbern(ilogit(alpha + beta*a[i]))
}

# Track variances
V_a <- sd_a^2
V_res <- res_sd^2
Vp0 <- sd_a^2 + res_sd^2
}"
writeLines(missingModgen,"./missingModgen.jags")

dat2 <- list(
n_ind=dim(d)[1],
n_d=n.dams,
n_s=n.sires,
z_obs=d$z2_obs,
dam=as.integer(as.factor(d$dam)),
sire=as.integer(as.factor(d$sire)),
s=d$W1
)

m2 <- jags.model(file="./missingModgen.jags",data=dat2,n.chains=2,n.adapt=1000,quiet=TRUE)
s2 <- jags.samples(model=m2,variable.names=c("mu","alpha","beta","V_a","V_res","Vp0","a"),n.iter=10000,thin=10,quie
```

The model convergence was checked for each run. Example output:

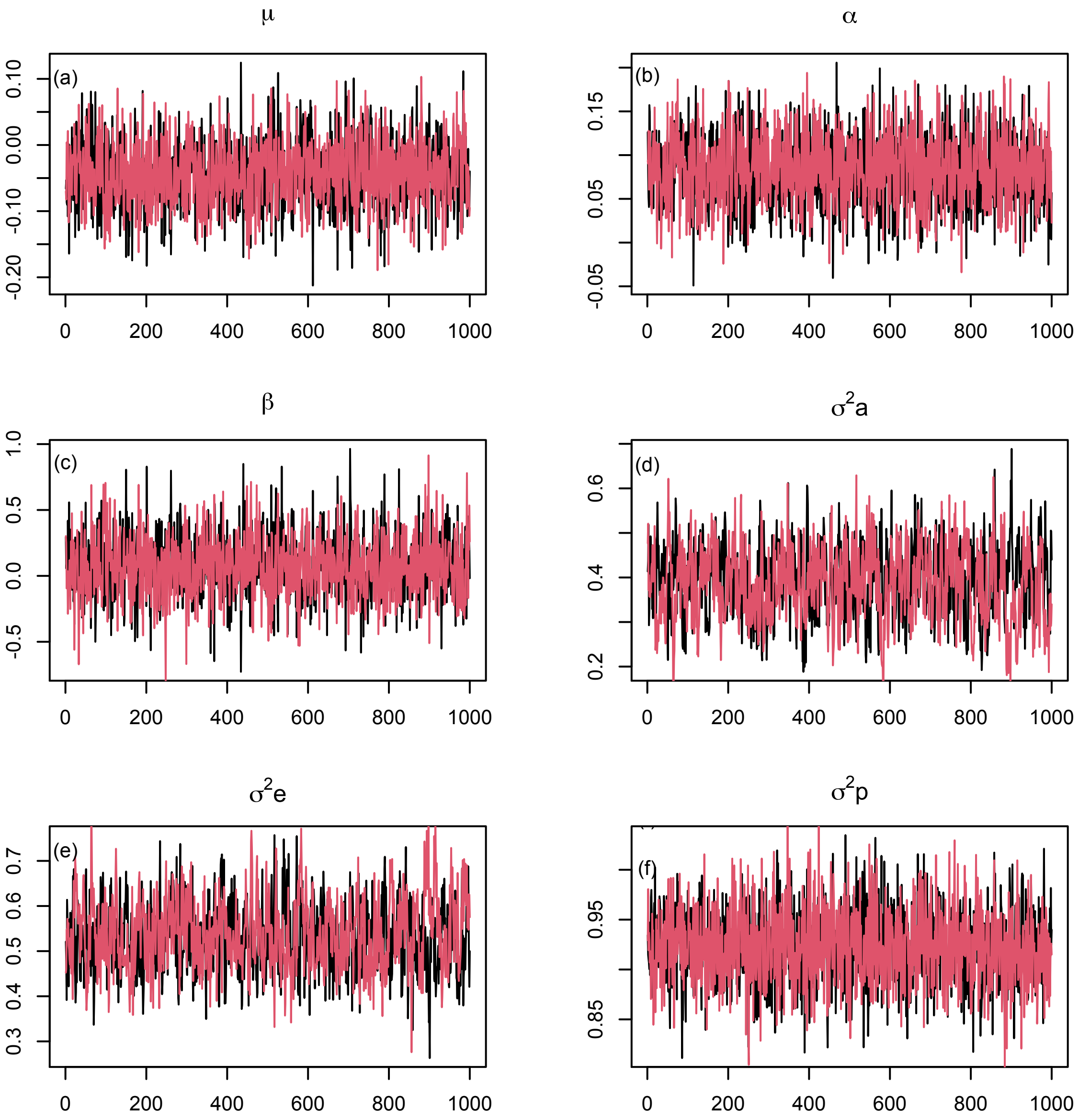

Figure 4: The convergence of the model parameters for an example analyses on simulated data where the strength of viability selection on trait 1 was 0.9 and the phenotypic covariance was 0.8.

##### When could we estimate phenotypic selection differentials without knowledge of the early-life trait?

Although we have said that the inference method can only pick up *genetic* selection differentials of trait 2, we can also obtain reasonable estimates of the phenotypic $S_{z_2}$ under certain scenarios where there is a high phenotypic correlation between the early-life trait(s) under viability selection and later-life trait(s) of interest. This is purely theoretical for many traits as it would be impossible to know the phenotypic covariance without the phenotypes of the early-life trait, and in that instance the episodes of selection method should be used. However, if you could justify that there is a strong phenotypic covariance with the later-life trait and whatever trait(s) are underlying prior viability selection, then this method would be possible. For example, in instances where body size traits are known to have a strong influence on fitness both early and late in life, and are also likely to have high a phenotypic correlation throughout life. The genetic relationship between those who died and the survivors is still required. With a full-sib design that could be knowing the nest-of-origin of chicks rather than a pedigree. The following is based upon the full-sib data as above.

**Analysis**: We ran mixed-models (R package `rjags`; R Core Team 2018; Plummer 2019) of phenotype (where $z$ is the trait we would measure in the second episode; i.e. $z_2$)

$$z_{ij} \sim N(\mu + b_j + e_{ij}), \tag{3}$$

with a random effect of genetic relationships (e.g., nest-of-origin or sibship; $b_j \sim N(0, \epsilon_j)$ where $j$ represents the family and $\epsilon$ is the residual) and a regression of prior survival ($W_a$ is survival after selection for $z_1$ in $y = a$) on later phenotypes ($z = z_2$)

$$W_{a,ij} \sim Bern(g^{-1}(\alpha + \beta z_{ij})). \tag{4}$$

These models were informed by an observation model for whether an individual was alive post-viability selection

$$z_{ij} \sim N(z_i^* + \eta), \tag{5}$$

where $z_i^*$ is the observed phenotype for $z_2$ in $y = b$ for individual $i$ with a high precision $\eta$.

Selection differentials were estimated from the logistic intercept ($\alpha_{z_2}$) and slope ($\beta_{z_2}$), trait means ($\mu_{z_2}$) and standard deviation ($\sigma_{z_2}$) obtained from these models in equations 14, 15 and 16 shown in the main manuscript.

```
# Jags simulation model using the observed later-life trait values,
# the knowledge of who survived/died
# and the family that these individuals belonged to
missingMod <- "model{
# Priors
fam_sd ~ dunif(0,3)
res_sd ~ dunif(0,3)
fam_tau <- 1/fam_sd^2
res_tau <- 1/res_sd^2
mu ~ dnorm(0,0.0001)
alpha ~ dnorm(0,0.0001)
beta ~ dnorm(0,0.0001)

# Random effect of family
for(j in 1:n_fam){
    b_fam[j] ~ dnorm(0,fam_tau)
}

# Simulation
for(i in 1:n_ind){
    # Phenotypes
    z[i] ~ dnorm(mu + b_fam[fam[i]], res_tau)
    # Observed phenotypes with little error
```

```
    z_obs[i] ~ dnorm(z[i], 1000)
    # Survival probability based on the fitness function
    s[i] ~ dbern(ilogit(alpha + beta*z[i]))
}

# Track variances
V_fam <- fam_sd^2
V_res <- res_sd^2
Vp0 <- fam_sd^2 + res_sd^2
}"
writeLines(missingMod,"./missingMod.jags")

# data required
dat <- list(
n_ind=dim(d)[1], # observed individuals
n_fam=n.sires, # number of families/nests/clonal lines
z_obs=d$z2_obs, # observations of the later-life phenotypes
fam=d$sire, # family individual is from
s=d$W1 # did the individual survive viability selection or not
)

m1 <- jags.model(file="./missingMod.jags",data=dat,n.chains=2,n.adapt=1000,quiet=TRUE)
s1 <- jags.samples(model=m1,variable.names=c("mu","alpha","beta","V_fam","V_res","Vp0","s"),
                   n.iter=100000,thin=100,quiet=TRUE)

# Composite function to calculate the prior viability selection differential from the
# logistic intercept and slope (a and b), trait mean and sd (m and s)
S_logistic <- function(a,b,m,s){
# get the sample range
    low <- m - 6*s; up <- m + 6*s;
# function to get absolute fitness
    fun <- function(z,a,b,m,s){(exp(a+b*z)/(1+exp(a+b*z)))*dnorm(z,m,s)}
# obtain absolute fitness
    muW <- integrate(f=fun,lower=low,upper=up,a=a,b=b,m=m,s=s)$value
# function to get covariance between absolute fitness and trait 2
    ffun <- function(z,a,b,m,s){(z-m)*(exp(a+b*z)/(1+exp(a+b*z)))*dnorm(z,m,s)}
# normalise to relative fitness
    return((integrate(f=ffun,lower=low,upper=up,a=a,b=b,m=m,s=s)$value-m*muW)/muW)
}

# Estimate the prior viability selection differential using parameters inferred from jags
post.S <- array(dim=1000)
for(i in 1:1000){
    post.S[i] <- S_logistic(s1$alpha[1,i,1],s1$beta[1,i,1],s1$mu[1,i,1],sqrt(s1$Vp0[1,i,1]))
}
mean(post.S)

# What should the prior viability selection differential have been?
# Equation 4 in the main manuscript
P0[1,2]/P0[2,2]*S_logistic(alpha,beta,0,P0[1,1])

S1_viab <- weighted.mean(d$z1,d$W1)-mean(d$z1) # viability selection on z1
S2_viab <- weighted.mean(d$z2,d$W1)-mean(d$z2) # effect of viability selection on z2

d2 <- subset(d,d$W1==1) # Just those individuals that survived
# Total selection of trait 2, but with missing fraction problem -- i.e. fecundity selection on z2
S2_fec <- weighted.mean(d2$z2,d2$W)-mean(d2$z2)
```

```r
# Total selection of trait 2, if we had perfect knowledge
S2_total <- weighted.mean(d$z2,d$W)-mean(d$z2)

# Covariance of z1, z2 and z2_obs
covar <- cov(d[,c("z1","z2","z2_obs")],use="pairwise.complete.obs")
```

When phenotypic covariance is high, we can actually recover the correct estimates of $S_{z_2,i}$ on a phenotypic level by running a jags model to obtain $\alpha_{z_2}$, $\beta_{z_2}$, $\mu_{z_2}$ and $\sigma_{z_2}$. These values can be used as input for the equations that integrate over fitness. When we do this we get a good estimate of $S_{z_2,i}$ at high phenotypic covariance (Fig 5a), but that we obtain overestimates at lower phenotypic covariances (Fig. 5b).

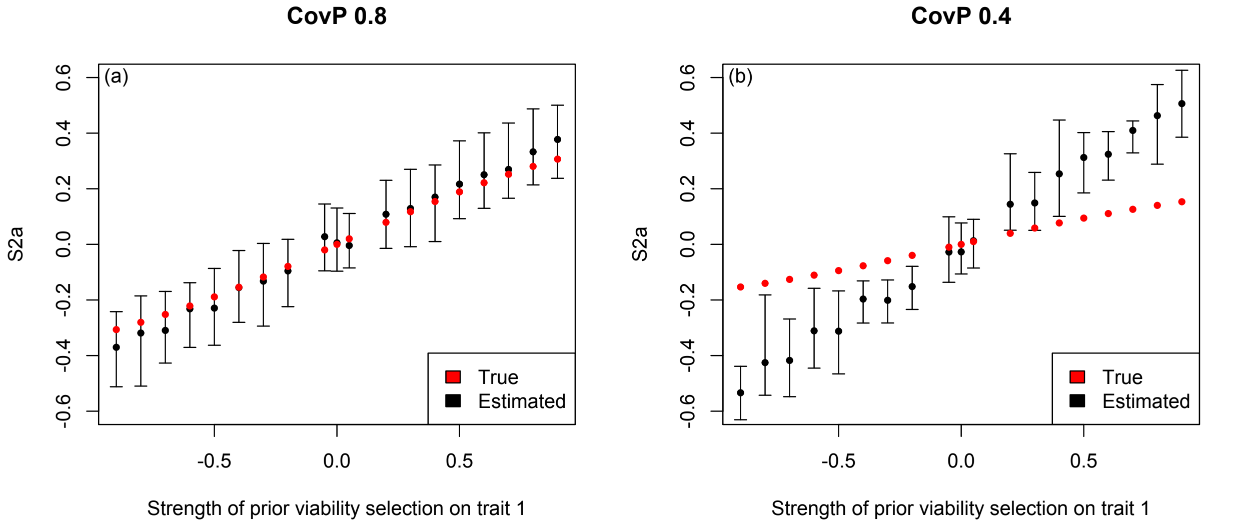

Figure 5: The true and inferred phenotypic selection differentials of trait 2 from simulated data of two traits expressed early (trait 1) and late (trait 2) in life that have (a) high phenotypic covariance ($\sigma(z_1, z_2) = 0.8$) and (b) weaker phenotypic covariance ($\sigma(z_1, z_2) = 0.4$). The true and inferred values are shown in red and black, respectively. The 95% confidence intervals around the estimated values are shown.
